# Nanobodies counteract the toxicity of an amyloidogenic light chain by stabilizing a partially open dimeric conformation

**DOI:** 10.1101/2023.08.28.555063

**Authors:** Broggini Luca, Barzago Monica Maria, Speranzini Valentina, Schulte Tim, Sonzini Federica, Giono Matteo, Romeo Margherita, Milani Paolo, Caminito Serena, Mazzini Giulia, Rognoni Paola, Merlini Giampaolo, Pappone Carlo, Anastasia Luigi, Nuvolone Mario, Palladini Giovanni, Diomede Luisa, Ricagno Stefano

## Abstract

Light chain amyloidosis (AL) is a systemic disease where fibrillar deposition of misfolded immunoglobulin light chains (LCs) severely affects organ function and results in poor prognosis for patients, especially when heart involvement is severe. Particularly relevant in this context is the cardiotoxicity exerted by still uncharacterized soluble LC species. Here, with the final goal of identifying alternative therapeutic strategies to tackle AL amyloidosis, we produced five llama-derived nanobodies (Nbs) specific against H3, a well-characterized amyloidogenic and cardiotoxic LC from an AL patient with severe cardiac involvement. We found that Nbs are specific and potent agents capable of abolishing H3 soluble toxicity in *C. elegans in vivo* model. Structural characterization of H3-Nb complexes revealed that the protective effect of Nbs is related to their ability to bind to the H3 V_L_ domain and stabilise an unexpected partially open LC dimer in which the two V_L_ domains no longer interact with each other. Thus, while identifying potent inhibitors of LC soluble toxicity, we also describe the first non-native structure of an amyloidogenic LC that may represent a crucial step in toxicity and aggregation mechanisms.

## INTRODUCTION

Systemic immunoglobulin light chain (AL) amyloidosis is a protein misfolding disease caused by the conversion of patient-specific immunoglobulin light chains (LCs) from their soluble functional states into highly organized amyloid fibrils (1). AL amyloidosis originates from an abnormal proliferation of a plasma cell clone that results in LCs overproduction and over-secretion in the bloodstream (2).

In AL amyloidosis, LCs misfold and assemble into amyloid fibrils that deposit in various organs leading to their dysfunction (1). More than 75% of AL patients have large fibrillar deposits in the heart, developing a rapidly progressive form of cardiomyopathy (3). Current therapies for managing AL amyloidosis rely primarily on eradicating the plasma clone to reduce the secretion of the misfolding-prone and toxic LCs (4). However, the treatment regimens required to eliminate the plasma cell clone are not always well tolerated by patients, especially when heart involvement is severe, highlighting the need for new therapeutic targets and strategies (5,6).

Besides the alterations caused by amyloid accumulation in the extracellular space, direct proteotoxicity exerted by prefibrillar soluble LC species is a further pathogenic factor (7–14). Indeed, clinical observations indicate that the reduction of circulating toxic LCs leads to rapid improvement in heart function and prognosis, despite no appreciable reduction in amyloid deposits (15,16). These observations have been recapitulated in established experimental models, such as human (14,17) and rodent (18–21) cardiac cells, zebrafish (22,23), and the nematode *C. elegans*. In particular, *C. elegans* was demonstrated as a robust model system to investigate the cardiotoxicity of amyloidogenic LCs (24,25). Indeed, the administration of cardiotropic LCs to worms alters severely the function of the pharynx, which is considered an “ancestral heart”. These alterations lead to mitochondrial injury and increased production of reactive oxygen species. Conversely, non-amyloidogenic LCs do not trigger significant toxic effects (24–26).

Over-expressed free LCs assemble as homo-dimers with a characteristic dimer interface (27). Each LC monomer comprises a N-terminal variable domain (V_L_) and a C-terminal constant domain (C_L_) (28) (Figure S1). The V_L_ domain spans about 110 amino acids and is characterized by high sequence variability, especially in the three hypervariable complementarity determining regions (CDRs) (Figure S1B). Conversely, the C_L_ domain is well conserved within the isotypes.

Even though the molecular mechanisms underlying LC cardiotoxicity remain largely unclear, several biochemical and biophysical traits were found to be typical of amyloidogenic LCs (27). In particular, strong evidences indicate that pathogenic LCs present a remarkable fold instability, increased dynamics and flexibility, increased susceptibility to protease cleavage, and ability to interact with metal ions (8,26,27,29–34). In addition, it has recently been reported that AL LCs exhibit a rather loose V_L_-V_L_ interaction, and this might explain their higher amyloidogenicity compared to non-AL LCs (35).

Morgan and co-workers have recently demonstrated that LC ligands which bind the native state impact amyloid fibril nucleation and elongation, stopping the amyloidogenicity cascade at its origin (33,36,37). On the same line, we have recently showed that introduction of conservative mutations into the V_L_ domain of a cardiotoxic LC, which leads to a stabilization of the fold, results in the abrogation of its soluble toxicity in *C. elegans* (8). Hence, our and other studies suggested LC fold stability as key to counteract AL amyloidosis pathology (38). We hypothesized that highly specific protein binders such as nanobodies (Nbs) could stabilize the native LC dimer structure. Nbs, or VHH antibodies, are monomeric antigen binding domains derived from camelid heavy-chain only antibodies (39). Despite being approximately one-tenth of the size of a conventional antibody, they retain specificity and affinity similar to conventional immunoglobins (40) but are easier to clone, express, and manipulate (41). Their small size (∼15 kDa), high water solubility, stability, specificity, ease of production, and robustness renders them ideal LCs stabilizers.

Here, we present five llama-derived Nbs directed to H3, an amyloidogenic LC belonging to IGLV1-44, from an AL patient with severe cardiac involvement (27). We demonstrated that the binding of the H3 by Nbs abolishes H3-driven toxicity on the pharynx of *C. elegans*. We showed that Nbs stabilize H3 and bind to their target with affinities in the nanomolar range. Finally, crystal structures of H3-Nb complexes show that Nb molecules form large interaction surfaces with monomeric V_L_ domains stabilising a non-native partially monomeric LC conformation.

Overall, our results show that by forming a stabilising complex between soluble LCs and a specific binder, it is possible to abrogate LC cardiotoxicity; moreover, we picture an unreported partially monomeric LC dimer representing a conformation likely specific for amyloidogenic and toxic LCs. Such conformation may be the one directly responsible for the observed soluble cardiotoxicity or at least an obligate step towards the formation of the toxic species.

## RESULTS

### Llama immunization and nanobodies generation

Cardiotoxic homo-dimeric LC H3 (Figure 1), produced recombinantly in *E. coli* and purified to homogeneity, was sent to NabGen Technology for llama immunization, yielding strong H3 binders after two panning rounds of phage-display coupled to ELISA. Five candidates, referred to as C2, C4, B5, C10 and H19, were selected based on the diversity of their CDR3 loops (Figure 1 and Table S1). Sequence alignment of the five Nbs revealed different CDRs and CDR3 regions with varying lengths between 14 and 17 residues (Figure 1A). After sub-cloning of the sequences for periplasmic expression in *E. coli*, Nbs were isolated by affinity and size-exclusion chromatography (SEC).

**Figure 1.**
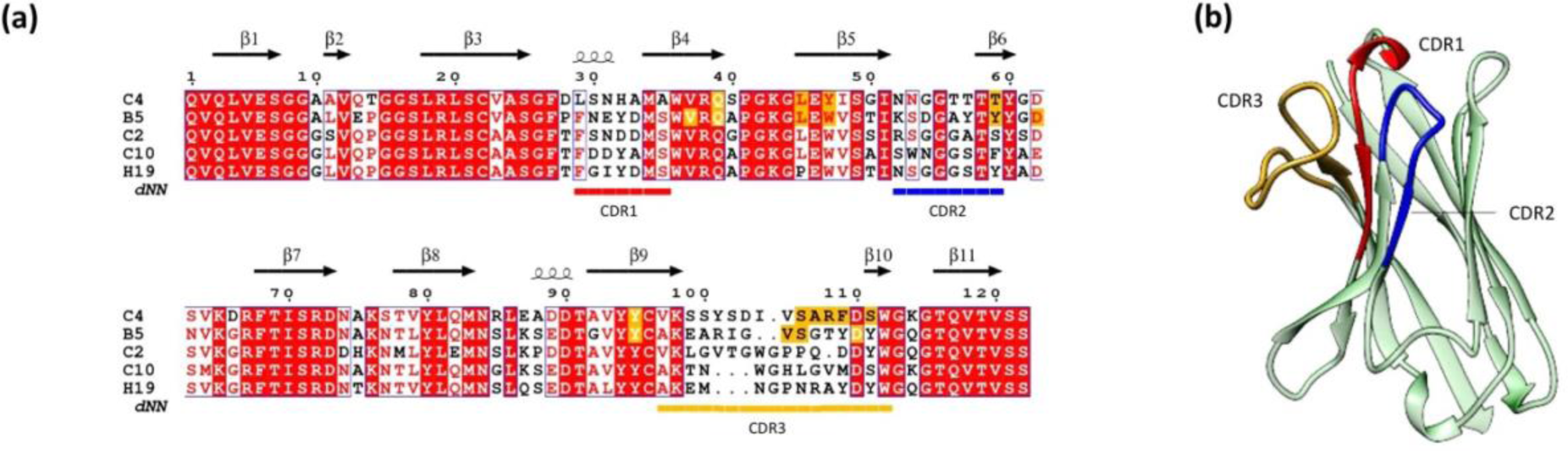
Features of Nbs sequences and fold. (a) Alignment of Nbs amino acid sequences. Sequence identity is indicated by a red box with white character, similarities within and across groups are indicated by red characters and blue frames, respectively. Secondary structure elements of the Nbs are shown above the sequence alignment and were extracted from the C4 structure. CDR1, CD2, and CDR3 are highlighted by red, blue, and gold bars, respectively, positioned below the sequence alignment. CDR regions were determined using the Integrated Nanobody Database for Immunoinformatics (http://research.naturalantibody.com/nanobodies). Residues involved in H3-Nb interaction are highlighted in orange. The alignment was visualized using ESPript 3.0 (41). (b) Nb fold is characterized by the typical immunoglobulin-like domain, consisting of a beta sandwich of 8 antiparallel β-strands arranged in two β-sheets with a Greek key topology. CDRs are highlighted in red (CDR1), blue (CDR2), and gold (CDR3).

### Nanobodies abolish H3 toxicity in *C. elegans* model

To assess the potential of the five Nbs to neutralize H3 toxicity, we compared the effect of SEC-isolated H3, Nbs and H3-Nb complexes on *C. elegans* pharyngeal function (Figure 2 and Figure S2) (8,24,26).

**Figure 2.**
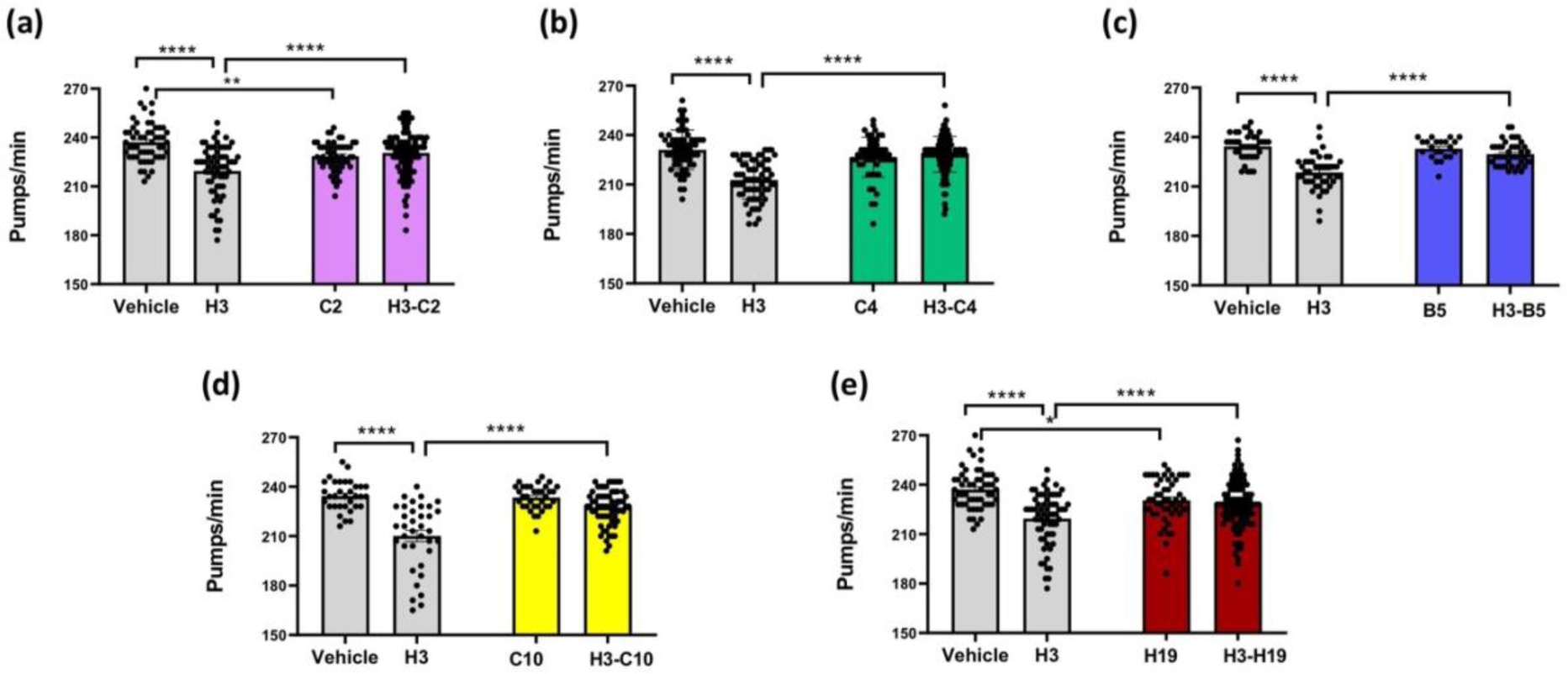
Effect of different Nbs on the toxicity caused by H3-LC protein in *C. elegans*. Worms were fed (100 worms/100 µL) with 100 µg/mL of H3, 100 µg/mL (a) C2, (b) C4, (c) B5, (d) C10, or (e) H19 Nb suspended in 10 mM phosphate buffered saline (PBS), pH 7.4 (Vehicle + Nb), or 100 µg/mL of a complex formed by each different Nb with H3 (H3-Nb dissolved in 10 mM PBS, pH 7.4). Control worms fed 10 mM PBS, pH 7.4 (100 worms/100 µL) (Vehicle). Pharyngeal pumping was determined 24 h after the administration by scoring the number of times the terminal bulb of the pharynx contracted over a 1-min interval (pumps/min). Each value is the mean ± SE (N=28-114). ****p<0.0001, ***p<0.001, **p<0.01 and *p<0.05, two-way ANOVA and Bonferroni’s *post hoc* test. Interaction: B5/H3=**** p<0.0001, C4/H3= **** p<0.0001, C2/H3= **** p<0.0001, H19/H3= ****p<0.0001 and C10/H3= ****p<0.0001, two-way ANOVA and Bonferroni’s *post hoc* test.

As reported previously (24), the pumping rate of H3-fed worms was significantly reduced compared to vehicle-fed ones (Figure 2). Conversely, administration of any H3-Nb complex did not alter significantly the pharyngeal activity compared to control (Figure 2). Interestingly, all H3-Nbs complexes had comparable effects, as they all prevented the pharyngeal dysfunction observed when H3 is administered alone (Figure 2). The administration of Nbs (C2 and H19) resulted in a slight, although significant, reduction of the *C. elegans* pharyngeal activity (Figure 2A and 2E). Nevertheless, administration of H3-C2 and H3-H19 complexes did not reveal any significant difference to controls (Figure 2A and 2E).

Lastly, to further prove the ability of Nbs to bind and limit H3 cardiotoxicity, we incubate H3 and monomeric Nb C4 for 30 min before co-administration to the worms (Figure S3). While dimeric H3 significantly affected the nematode pharyngeal activity, H3 co-administered with C4 had no effect, as no significant reduction in the pumping rate was observed (Figure S3).

Overall, all five tested Nbs effectively abolished the soluble toxicity of H3 administered in the *C. elegans* model system.

### Nanobodies bind with nanomolar affinity to H3 and form stable complexes

To determine thermodynamic and kinetic parameters of H3-Nb binding, we applied isothermal titration calorimetry (ITC), and bio-layer interferometry (BLI), respectively (Table 1). Injection of Nbs into H3 yielded ITC thermogram profiles with initial exothermic peaks that returned to baseline at the end of the titration, indicating binding saturation (Figure S4). Control titrations of Nb into buffer did not yield any signal above background (Figure S4). The derived binding isotherms were fitted with a one-site binding model, revealing strong binding affinities in the nanomolar range for all tested H3-Nb pairs, (Figure 3A and Table 1). In particular, Nbs C4 and C10 bound with K_D_ values of 17 ± 10 nM and 31 ± 23 nM, respectively. Binding of C2, B5 and H19 was somehow weaker, with affinities of 402 ± 85 nM, 350 ± 79 nM and 176 ± 70 nM respectively, although still within the nanomolar range. ITC also revealed exothermic enthalpy (ΔH) values ranging between −20.4 ± 0.7 kcal/mol and −10.9 ± 0.4 kcal/mol (Table 1). ITC experiments point to a 1:1 stoichiometry of the H3-Nb complex (Figure 3A).

**Figure 3.**
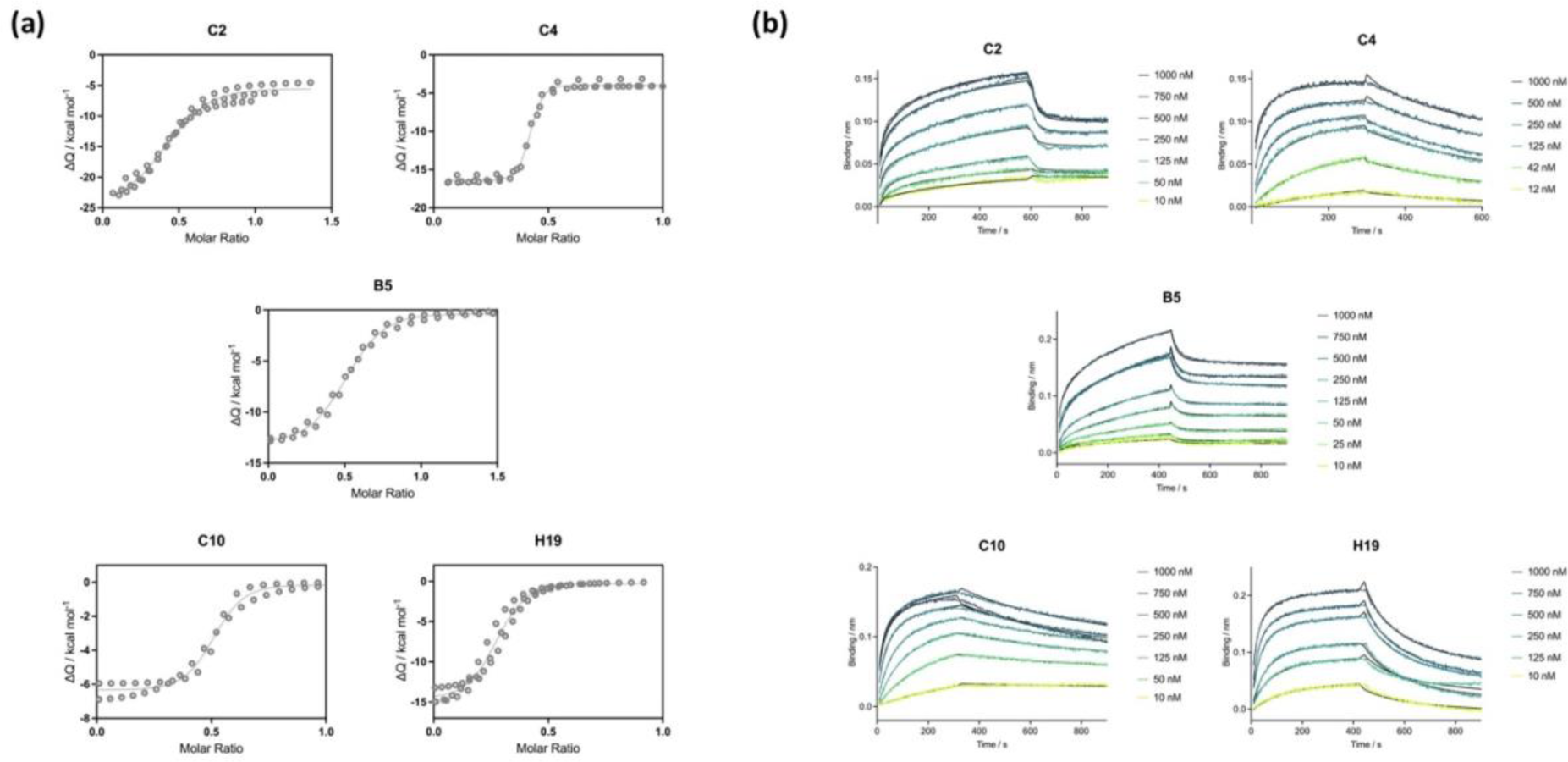
H3-Nb binding characterization. (a) ITC binding isotherms obtained from at least two experiments were fit globally applying a 1:1 binding model (molar ratio refers to LC dimer). Associated thermograms are shown in Figure S3. (b) BLI sensorgrams revealed H3 concentration-dependent binding to surface-immobilized Nbs. Sensorgrams are color-coded according to the H3 concentration. Bayesian 1:1 binding models are shown as black solid lines.

**Table 1.** Kinetic and thermodynamic parameters of H3-Nb binding.

| Complex | ITC |  | BLI |  |  |  |
| --- | --- | --- | --- | --- | --- | --- |
| | $K_D$ / nM | $\Delta H$ / kcal/mol | $K_D$ / nM | $K_{off}$ / $s^{-1}$ | $K_{on}$ / $M^{-1} s^{-1}$ | $K_D$ steady-state / nM |
| H3-C2 | $402 \pm 85$ | $-(20.4 \pm 0.7)$ | 848 | $4.33E-02$ | $5.11E+03$ | 167 |
| H3-C4 | $17 \pm 10$ | $-(12.6 \pm 0.3)$ | 12 | $1.52E-03$ | $1.24E+04$ | 81 |
| H3-B5 | $350 \pm 79$ | $-(15.3 \pm 0.6)$ | 540 | $4.33E-02$ | $8.02E+03$ | 255 |
| H3-C10 | $31 \pm 23$ | $-(10.9 \pm 0.4)$ | 35 | $1.18E-03$ | $3.34E+03$ | 84 |
| H3-H19 | $176 \pm 70$ | $-(14.8 \pm 1.8)$ | 316 | $8.11E-03$ | $2.57E+03$ | 121 |

BLI measurements confirmed the strong-affinity binding of Nbs to H3 (Figure S5 and Table 1). Dipping of surface-immobilized Nbs into H3 yielded binding curves with maximal response levels reaching between 0.15 and 0.2 nm at the highest tested H3 concentrations (Figure 3B). Global Bayesian fits applying a 1:1 binding model to the datasets revealed binding affinities close to the ones obtained by ITC (Table 1). Similarly, the derived semi-log concentration– response curves revealed sigmoidal line-shapes with fitted apparent K_D_ values in the same nanomolar range (Figure S5). Nbs C4 and C10 with the highest binding strength in ITC also exhibit the slowest dissociation rates in BLI (Table 1).

Then, to evaluate the effect of Nb complexation on the stability of dimeric H3 native fold, we monitored thermal unfolding by intrinsic fluorescence (Figure S6). All H3-Nb complexes showed a high degree of cooperativity during thermal unfolding, well described by a two-state model, except for H3-C10, which presented an unfolding profile likely affected by an aggregation event (Figure S6). The melting temperatures (T_m_) values indicate a general, though moderate, increase in thermal stability for all the complexes compared to isolated H3 (Table 2). The largest stabilization effects were observed for H3-H19, that showed a T_m_ increment of and 4.0 °C compared to isolated H3. Conversely, H3-C2 transition midpoint was found at 55.5 ± 1.1 °C, 2.2°C lower than H3 (57.2 ± 0.4 °C).

**Table 2.** Melting temperature values for H3-Nb complexes. *H3-C10 presents a peculiar unfolding profile (Figure S7) due to protein aggregation at high temperatures that affects fluorescence signals. The T_m_ was derived assuming the cooperative behaviour observed for the other complexes.

| Complex | Melting temperature / °C |
| --- | --- |
| H3 | $57.2 \pm 0.4$ |
| H3-C2 | $55.5 \pm 1.1$ |
| H3-C4 | $59.6 \pm 0.5$ |
| H3-B5 | $59.0 \pm 0.2$ |
| H3-C10 | $59.4 \pm 0.2^*$ |
| H3-H19 | $61.2 \pm 0.6$ |

Overall, these data demonstrated the formation of stable H3-Nb complexes with K_D_ values in the nanomolar range. Notably, our analyses revealed Nbs C4 and C10 as best binders with strongest affinity and slowest dissociation from H3. Thermal unfolding measurements indicate a moderate binding-induced stabilization of H3 in complex with these Nbs.

### Nbs C4 and B5 bind to the same structural epitope of H3 V_L_ domain

To understand the structural basis underlying the neutralization effect of Nbs on H3 toxicity, we determined the crystal structures of two H3-Nb complexes, namely H3-C4 and H3-B5, at 2.0 Å and 3.2 Å resolution, respectively (Figure 4 and Table 3). H3-C4 exhibited a dimer interface with a two-fold axis of symmetry, resulting in the presence of a single H3-C4 pair (one LC molecule + one Nb molecule) in the asymmetric unit. In contrast, the asymmetric unit of H3-B5 consisted of the H3 dimer decorated with two B5 molecules. Both structures clearly showed that Nbs C4 and B5 bind identically to H3, with the H3-Nb single pair conformation perfectly superimposable (RMSD value over 330 Cα of 1.18 Å) (Figure 4C). Furthermore, in both cases, the H3 dimer is found interacting with two Nb moieties, confirming the stoichiometry inferred from ITC data (Figure 4A).

**Figure 4.**
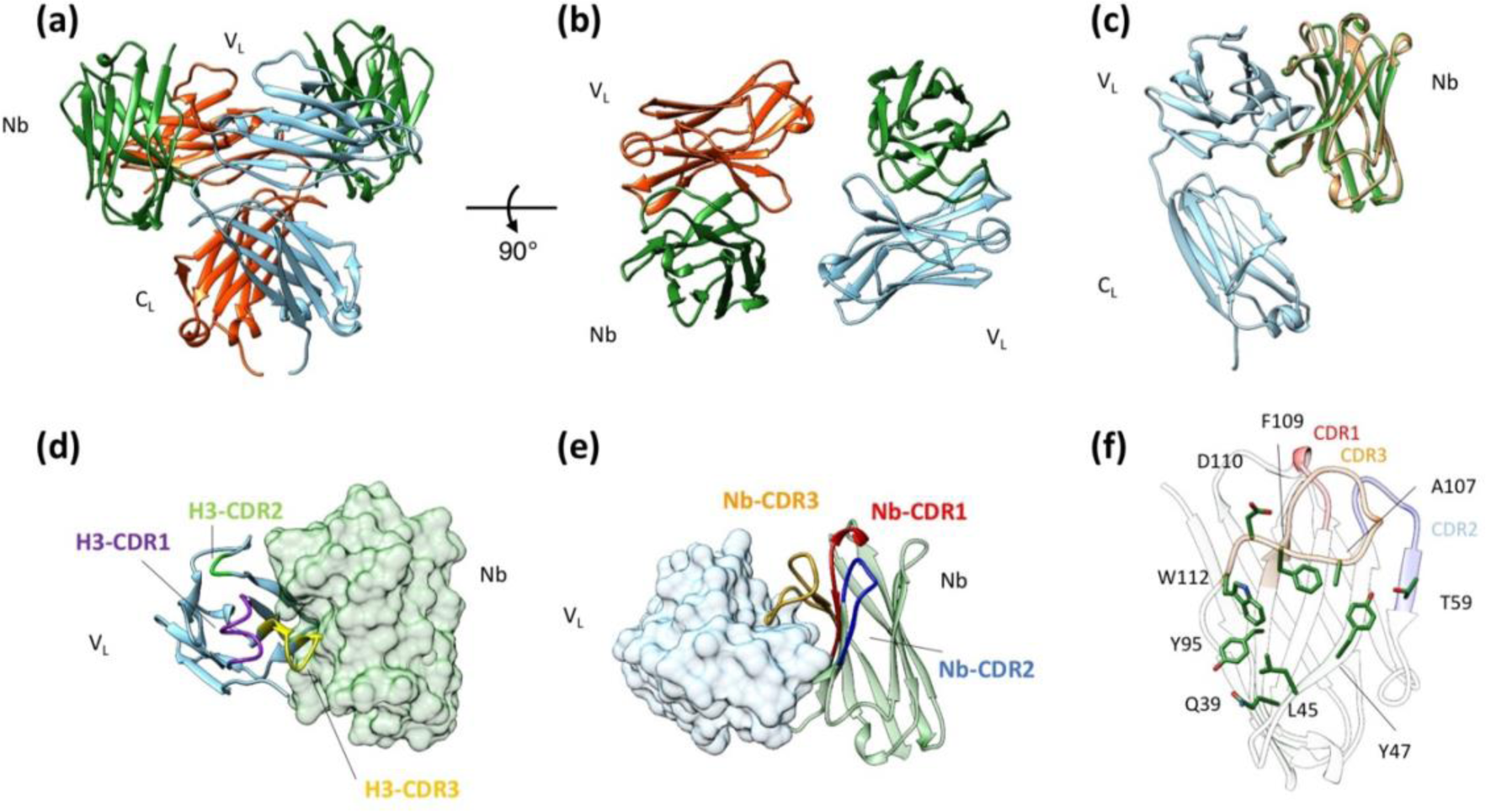
Nbs bind to the V_L_ of H3. (a) 3D architecture of the H3 dimer (monomers in cyan and red) in complex to two C4 nanobodies (green), each decorating the same structural epitope of H3-V_L_. (b) Top view of H3-C4 interaction, with the two Nbs each binding a single V_L_ domain. (c) Nbs C4 (green) and B5 (orange) bind in an equal manner to H3 (cyan). The RMSD value between monomeric H3-C4 and monomeric H3-B5 was 1.18 over 330 Cα. (d) C4, represented as green surface, specifically target CDR3 (gold) and the region joining CDR1 (violet) and CDR2 (green) of H3 V_L_ (cyan). (e) The primary contacts between H3 V_L_ (cyan, surface) and C4 (green) are mediated through C4 CDR3 (orange), CDR2 (blue), and the β-sheet connecting CDR1 (red) and CDR2 (blue). (f) C4 (green) fold with residues involved in H3 V_L_ binding represented as sticks. CDRs are reported and highlighted in red (CDR1), blue (CDR2), and orange (CDR3).

**Table 3.** Data collection and refinement statistics for the crystal structures of the H3-C4, and H3-B5 complexes.

| Structure | H3-C4 | H3-B5 |
| --- | --- | --- |
| PDB ID | 8P88 | 8P89 |
| Beam Line | ID30B | ID30B |
| Space group | C 2 2 21 | I 41 2 2 |
| Unit cell constants (Å, °) | a = 73.6, b = 121.9, c = 88.7;<br>$\alpha = 90, \beta = 90, \gamma = 90$ | a = 145.2, b = 145.2, c = 245.2<br>$\alpha = 90, \beta = 90, \gamma = 90$ |
| Resolution (Å) | 50.24 - 2.02<br>(2.16 - 2.02) | 124.9 – 3.18<br>(3.55 - 3.18) |
| CC (1/2) | 0.99 (0.65) | 0.99 (0.77) |
| I/σI | 18.2 (1.7) | 6.7 (1.5) |
| Completeness (spherical) | 100 (22.1) | 99.6 (11.6) |
| Completeness (ellipsoidal) | 100 (56.6) | 99.6 (79.7) |
| Multiplicity | 11.1 (12.7) | 11.5 (15.3) |
| Unique reflections | 21041 | 14056 |
| Refinement |  |  |
| R<sub>work</sub> (%) | 22.18 | 23.0 |
| R<sub>free</sub> (%) | 23.34 | 27.8 |
| Number of atoms | 2752 | 4867 |
| Protein | 2531 | 4591 |
| Waters | 219 | 87 |
| Heteroatoms | 2 | 15 |
| Ramachandran plot, *n* (%) |  |  |
| Most favoured region | 294 (97.58%) | 624 (96.0%) |
| Allowed region | 7 (2.42%) | 26 (4.0%) |
| Outliers | 0 (0.0%) | 0 (0.0%) |

The structural analysis showed that CDR2, CDR3, and β-sheet connecting CDR1 and CDR2 of the Nb directly interact with the CDR3 and CDR1-CDR2 joining region of H3 V_L_ (Figure 4D and 4E). The H3 V_L_-Nb is mainly driven by hydrophobic interactions, as the interface between the two proteins is mostly composed of non-polar or slightly polar amino acids (Figure S7 and Table S2). A further important contribution to the stabilization of the complex is the presence of backbone and side chain hydrogen bonds and ionic interactions (Figure S7 and Table S3). Interestingly, the structural analysis also revealed that the two Nbs share key residues to optimize the binding to the target. Indeed, in the two complexes, Nb residues Gln39, Tyr95, and Leu45 establish crucial H-bonds (Gln39) and hydrophobic interactions (Tyr95 and Leu45) stabilizing H3 V_L_-Nb interface (Figure 4F and Figure S7). Similarly, the two Nbs presented common residues responsible to tighten the interaction with H3 V_L_ (Figure S8). In particular, in C4, the CDR2 residue Thr59 establishes a side chain H-bond with H3 V_L_, while residues Ala107, Phe109, Asp110, and Trp112 on the CDR3 loop make backbone H-bond interactions and/or consolidate the hydrophobic network at the interface with H3 V_L_ (Figure 4F and Figure S7 and S8). The same is true for Nb B5, in which, despite some sequence variabilities, the interaction network is similar to that of C4 as the chemistry of the residues involved is largely conserved (Figure S7 and S8 and Tables S2 and S3).

These observations prompted us to speculate on the binding mechanism of the remaining Nbs (C2, C10, and H19). Indeed, both sequence analyses and model predictions by Alphafold2 showed that the putative binding regions of C2, C10, and H19 are similar to the ones of the C4 and B5, both in terms of surface engaged for the binding and in terms of types of interactions, as the interface is mainly composed of hydrophobic or slightly polar residues (Figure 1A and Figure S9).

Overall, in both crystal structures, the Nbs engage almost identical structural epitopes of H3, comprising mostly the highly variable CDRs. Nbs bind their target in similar orientations, interacting through both CDR and framework regions.

### Nbs binding stabilises a partially open H3 dimeric conformation

Surprisingly, compared to the canonical closed LC dimer structure (27), H3 adopts an ‘open’ dimer conformation in complex with the Nbs. H3 undergoes a large conformational change upon complex formation, whereby the two V_L_ domains form significant interaction interfaces only with Nbs molecules. Such interactions totally replace the V_L_-V_L_ dimeric interface present in the H3 structure and which has been observed in all crystal structures of LC dimers so far reported (Figure 5A and 5C) (27,36,42,43). In the crystal structures of isolated LC dimers, monomers are not superposable because they display different elbow angles between the V_L_ and C_L_ domains (Figure 5A and 5C) (27,36,42,43). On the contrary, the binding of the Nb molecules causes the rotation of one V_L_ domain by approximately 108° (Figure 5B and 5D). As a result, in the H3-Nb complexes the two H3 monomers have identical conformations, corresponding to the most bent conformation found in the structures of isolated LC dimers, including H3 (27,42). Consequently, Nb molecules stabilise a partially open conformation of H3 where C_L_ domains form the standard dimeric interface while V_L_ domains are virtually monomeric (Figure 5B and 5D).

**Figure 5.**
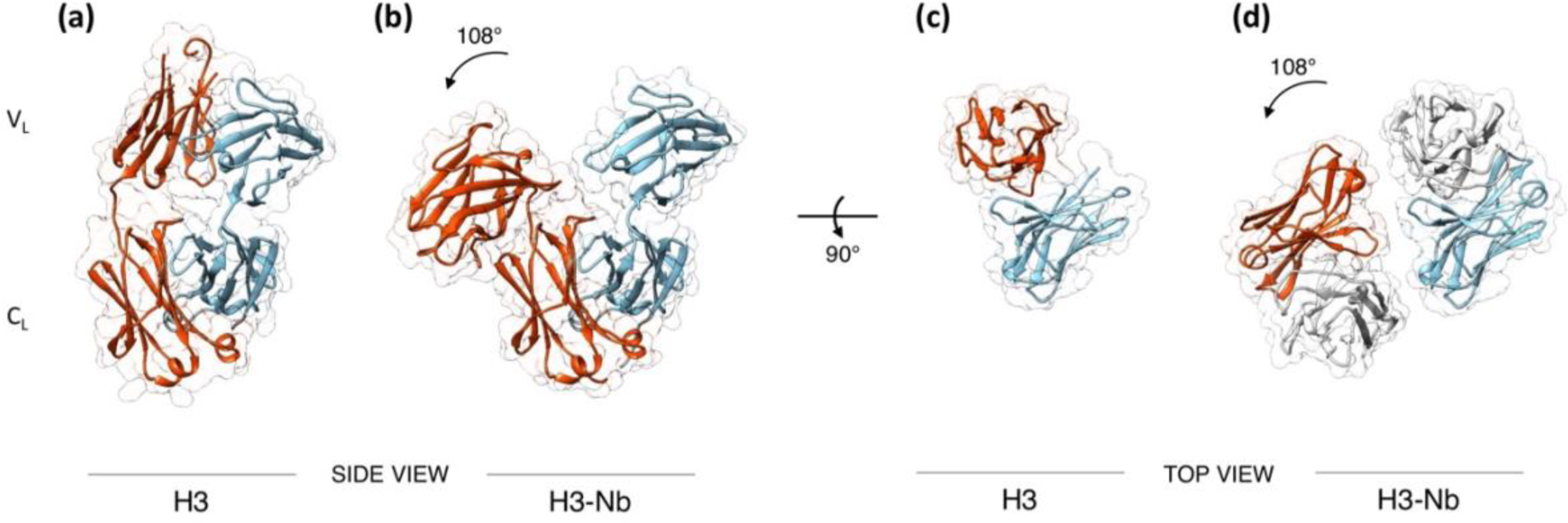
Nb binding stabilizes V_L_ domains in a partially open conformation. (a-b) Side-view comparison between conformations of (a) dimeric H3 (pdb 5MTL) and (b) H3-Nb. In the H3-Nbs complexes, the two V_L_ domains are separated from each other and are no longer at interaction distance. For clarity, Nbs are not shown. (c-d) Top-view comparison of the reciprocal orientation of V_L_ domains before (c) and after (d) Nb (in grey) loading. Each Nb interacts with a V_L_ domain and disrupts V_L_-V_L_ interactions. For comparisons, the two conformations were aligned on the C_L_ domains, that are not affected by Nb loading and retain the same reciprocal orientation.

To rationalise the effect of Nb binding on H3 architecture, we compared the interaction surfaces between V_L_-V_L_ in H3 dimer and H3-Nb complexes. Nbs binding closely resembles the orientation of the second V_L_ domain in the closed LC dimers. However, in native H3, the V_L_-V_L_ interface area measures 319.4 Å^2^, whereas the H3-Nb complexes exhibit more extensive interaction surfaces (794.9 Å^2^ and 732.0 Å^2^ for H3-C4 and H3-B5, respectively). These extended areas are also stabilized by a greater number of contacts, either hydrophobic and ionic (Figure 6A and 6B and Tables S2 and S3). Notably, while the H3 V_L_-V_L_ interface is solely composed by residues that fall in framework regions, in H3-Nb complexes H3 CDR residues are directly involved in the stabilization of the interaction. Indeed, in H3 native dimer, the stability of the V_L_-V_L_ interface is supported by specific framework residues, namely Phe101, Phe88, Ala44, Pro45, Tyr37, and Gln39 (Figure S7 and Tables S2 and S3). In H3-Nb structures, additional H3 residues play a role in the interaction. These include Lys46, His50, and Asn35 from the framework regions, as well as Trp92, Leu96, Asn97, and Val99 from the CDRs (Figure S7 and Tables S2 and S3). Furthermore, the higher number of H-bond interactions in H3-Nb structures suggests a tighter and more stable protein-protein interface (Figure S7 and Table S3). Interestingly, the stabilising interaction observed in the H3 dimer between the Gln39 residues belonging to the two monomers is replaced by the interaction with Gln39 present in all Nbs (Figure S7 and Table S3). The increased stability of the H3-Nb complexes was further confirmed by solvation free energy (ΔG_int_) calculations, revealing values of −15.5 kcal/mol, −36.0 kcal/mol and −42.4 kcal/mol for the V_L_-V_L_, V_L_-C4, and V_L_-B5 interfaces, respectively.

**Figure 6.**
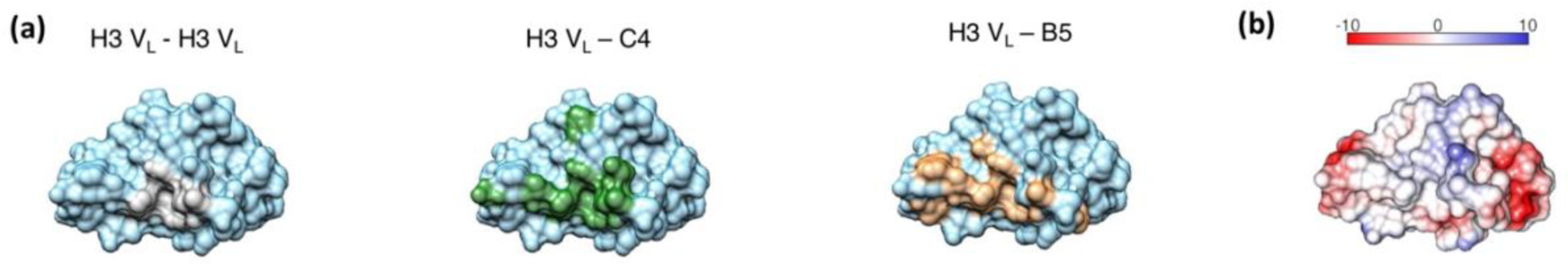
H3-Nb interfaces are larger than V_L_-V_L_ in dimeric H3. (a) Monomeric H3 V_L_ represented as surface with H3 V_L_ residues involved in V_L_-V_L_ interface highlighted in grey, V_L_-C4 in green, and V_L_-B5 in orange, respectively. (b) H3 V_L_ represented as surface and color-coded by electrostatic potential ranging from negative (red) to positive (blue) (scale in kcal/mol·e).

## DISCUSSION

Despite recent advances in diagnosis and treatment, the prognosis for AL patients with severe heart involvement remains very poor with a life expectancy of months (5). In such patients, the cardiotoxicity exerted by circulating soluble species is particularly important in impairing heart performance (7,13,15). To date, the molecular LC species responsible for such cardiotoxicity remain to be identified; however, mutations stabilising the native fold of a toxic LC result into a non-toxic variant (8). Moreover, it has been suggested that small molecule ligands acting as kinetic stabilisers of the LC dimeric native fold may prevent all aberrant downstream processes such amyloid aggregation and the formation of cardiotoxic species (37,38).

To experimentally verify if the use of highly specific molecular LC stabilizers could lead to the reduction of soluble LC toxicity, here we produced and characterised five llama-derived Nbs raised against H3, a λ LC responsible for severe cardiotoxicity in an AL patient (27).

The Nbs here presented are highly effective and potent H3 binders with nanomolar K_D_ values (Figure 3 and Table 1). Crucially, we found the Nbs to abrogate completely the soluble toxicity of H3 in *C. elegans*-based functional assays (Figure 2 and Figure S2). Notably, crystal structures revealed two Nbs to recognize the same structural epitope on the V_L_ domain (Figure 4C and Figure S9), comprising hypervariable segments of H3 V_L_ (Figure 4 and Figure 1B). AI-based modelling predicted similar binding modes for the remaining Nbs, in line with their comparable neutralizing effects in toxicity assays.

The structural analysis revealed that Nbs C4 and B5 primarily bind to the same structural epitope comprising hypervariable segments of H3 V_L_ domain (Figure 4 and Figure 1B). Crucially, this structural epitope is only accessible in the open dimer conformation of H3 (Figure 5B), but not the canonical ‘closed’ LC dimer structure (Figure 5A). It follows that these regions are accessible and were capable to trigger the formation of specific Nbs in the immunized llama. Thus, our data are directly showing that a partially dimeric open conformation of H3 - a conformation where the two V_L_ domains are virtually monomeric (Figure 5B and 5D) - exists and is significantly populated in solution. In other words, the Nbs binding epitope demonstrates that the weak V_L_-V_L_ interface present in H3 results in a dynamic equilibrium between a fully dimeric native conformation (Figure 5A and 5C) and a partially dimeric assembly, where the two V_L_ domains are effectively monomeric as seen in the H3-Nb crystal structures (Figure 5B and 5D). Interestingly, in such conformation, the V_L_ domains may have the same low stability and high aggregation propensity observed in isolated V_L_ domains (34,44–46): indeed in such open conformation, the V_L_ domains do not form any stabilizing intermolecular interactions. This lack of interactions makes them more susceptible to partial or complete unfolding, creating an ideal scenario for misfolding and the formation of non-native interactions with other V_L_ domains. In keeping with this model, the two main regions becoming solvent exposed upon V_L_ monomerization are largely hydrophobic and predicted to have a high aggregation propensity (Figure 6B and Figure S10). As recent Cryo-EM structures of fibrillar LCs show (47–50), the V_L_ domain undergoes a complete structural reorganisation from the native to the amyloid structure: it is intriguing to speculate that the partially open dimer presented in this work may represent the first step along the LC amyloid aggregation pathway.

The existence of this partially open dimer captured in H3-Nb structures prompts us to speculate how the Nbs may abolish the soluble toxicity effect of H3. One possibility is that this partially open dimer is the toxic species and the toxic effects observed *in vivo* are due to the exposure of hydrophobic clusters exposed by V_L_ domains once the V_L_-V_L_ interface is lost (Figure 6B and Figure S10). Such clusters may aberrantly interact with cellular components and membranes. In this case, the detoxification by Nbs is due to the protection and hiding of such regions. Another possibility is that Nbs stabilize this unstable open dimer, preventing further V_L_ unfolding and the formation of yet-to-identify highly hydrophobic oligomers directly responsible for the cardiotoxity as observed in other systems (51,52). While our data cannot discern between these two models, they strongly suggest that the open dimer here reported represents a species associated with soluble cardiotoxicity of H3.

Even though here we focused our characterisation on one cardiotoxic LC, our findings, in agreement with recent data, suggest that the presence of a partially open dimeric conformation may be a biophysical trait common to toxic/amyloidogenic LCs. Indeed, several evidences showed that increased dynamics, increased flexibility and exposure of aggregation-prone regions in amyloidogenic LCs are all crucial drivers of LC aggregation (27,53–57). In particular, Rennella et al. postulated that a distinctive property of aggregation-prone LCs is their loose V_L_-V_L_ interface, that results in increased susceptibility to misfolding and eventually aggregation (35). Similarly, a recent work by Rottenaicher and co-workers further emphasizes the correlation between labile dimer interface and propensity to misfold and aggregate in amyloidogenic LCs (58). On the same line, Gursky et al. recently showed that aggregation hotspots in LC sequence combined with the increased dynamics/exposure of such regions in isolated V_L_ domains trigger amyloid formation of κ LCs (59). This partially open dimer may account for the ability of full-length LCs to aggregate and be part of *in vivo* amyloid deposits (60–62).

In summary, high-affine and specific Nbs raised against circulating LCs counteract their biophysical instability responsible for the life-threatening cardiotoxicity. Thus, Nbs represent promising tools to mitigate the toxicity exerted by cardiotropic LCs in patients affected by AL amyloidosis with severe cardiac involvement. Such LC-stabilizing Nbs could complement existing clonal plasma cell eradication therapies for AL to reduce symptoms and increase the lifespan of patients.

## MATERIAL AND METHODS

### LCs purification

Recombinant LCs were produced according to Oberti et al. (27). Briefly, heterologous proteins, produced in the cytoplasm as inclusion bodies, were retrieved and subjected to a renaturation procedure, followed by purification through ion exchange and size exclusion chromatography. Recombinant LCs were biochemically characterized by Tycho and circular dichroism analyses to verify homogeneity and correct folding. Gel filtration analysis indicates that all LCs used in this work were dimeric in solution (Figure S1).

### Nb generation and purification

Immunization of a single llama was achieved with three subcutaneously injections at three-week interval of 0.5 mg of purified H3 in 10 mM Hepes pH 7.5, 150 mM NaCl. Lymphocytes were isolated from blood samples obtained 5 days after the last immunization. The cDNA was synthesized from purified total RNA by reverse transcription and was used as a template for PCR amplification to amplify the sequences corresponding to the variable domains of the heavy-chain antibodies. PCR fragments were then cloned into the phagemid vector pHEN4 to create a Nb phage-display library (63). The selection and screening of Nbs were performed as described previously (64). Three rounds of panning resulted in the isolation of specific binders. After sequence analysis, five different positive clones (namely C2, C4, B5, C10, and H19) were chosen. All the steps were performed by the NabGen Technology platform (https://nabgen.org/).

The Nbs sequences were cloned for expression in the pET28a+ plasmid with a pelB signal peptide for periplasmic expression at the N-terminus and a C-terminal 6xHis tag. The plasmid was used to transform BL21 cells. Cells were grown in Terrific Broth supplemented with 0.1% glucose and 25 µg/mL kanamycin. Expression was induced with 0.5 mM IPTG at OD_550_ of 0.6-0.8; after induction, cells were grown overnight at 18°C. Nbs were retrieved from the periplasm by osmotic shock: cells were resuspended in TES buffer (200 mM Tris-HCl, 0.5 mM EDTA, 500 mM sucrose, pH 8.0) and incubated for 1 h on ice. Lysis was obtained by addition of equal volume of TES buffer diluted 1:4 into water. The suspension was kept on ice for 2 h and then centrifuge at 4°C for 40 min at 20000 x *g*. After clarification, the supernatant was loaded onto a 5 mL Ni-NTA column pre-equilibrated in 50 mM Tris-HCl, 300 mM NaCl, 10% (v/v) glycerol, 15 mM imidazole, pH 8.0. Nb was eluted with a 2-step gradient at 50 mM and 250 mM imidazole and further purified through size exclusion chromatography using a Superdex 200 increase 10/600 column.

### H3-Nb complex formation

H3-Nb complexes were obtained by incubating dimeric H3 and monomeric Nb in a 1:2.2 molar ratio at 4°C for 16 h. Eventually, the mix was purified through SEC using a Superdex 200 increase 10/600 column equilibrated in 50 mM Hepes pH 8.0.0, 150 mM NaCl.

### Analytical size exclusion chromatography

Analytical SEC was performed using a Superdex 200 increase 10/600 column operated at 4°C by an Akta purifying system. Samples were injected onto the column extensively equilibrated with 50 mM Hepes pH 8.0.0, 150 mM NaCl. Runs were imported in GraphPad Prism 9.0 software (CA, USA) for data normalization, visualization and graph generation.

#### Effect of Nbs on the toxicity of H3 in *C. elegans*

Bristol N2 strain was obtained from the Caenorhabditis elegans Genetic Center (CGC, University of Minnesota, Minneapolis, MN, USA) and propagated at 20 °C on solid Nematode Growth Medium (NGM) seeded with *E. coli* OP50 (CGC) for food. The protective effect of six different NBs on the pharyngeal dysfunction caused by the administration to worms of H3 was investigated as already described (8,24,26). Briefly, worms (100 worms/100 µL) were incubated with 100 µg/mL H3, 100 µg/mL B5, C3, C4, C2 H19 or C10 Nbs in 10 mM phosphate buffered saline (PBS), pH 7.4, or 100 µg/mL of a complex formed by each different Nb with H3 in 10 mM PBS, pH 7.4. Control worms were administered with 10 mM PBS, pH 7.4 (100 worms/100 µL). After 2 h of incubation on orbital shaking, worms were transferred onto NGM plates seeded with OP50 *E. coli* and the pharyngeal pumping rate was scored 24 h later by counting the number of times the terminal bulb of the pharynx contracted over a 1-min interval. In same experiments, 100 µg/mL H3 was incubated for 30 min at room temperature with 100 µg/mL C4 Nb before the administration to worms (100 worms/100 µL). C4 alone (100 µg/mL) and H3 alone (100 µg/mL) were administered in the same experimental conditions and 10 mM PBS, pH 7.4, was administered as negative control. After 2h of incubation on orbital shaking nematodes were transferred to NGM plates seeded with fresh OP50 *E. coli* and the pumping rate was scored after 24h as previously described (24).

### Thermal unfolding ramps

Thermal unfolding experiments were performed using a Tycho NT.6 device following the changes in the intrinsic fluorescence detected at both 350 nm and 330 nm. Temperature ramps were performed in 50 mM Hepes pH 8.0, 150 mM NaCl from 35°C to 95°C. Melting temperature is defined as the temperature at which the folding-to-unfolding transition occurs and is the maximum or minimum of the 350 nm / 330 nm ratio curve first derivative. Each experiment was performed in triplicate. Raw data were imported in GraphPad Prism 9.0 software (CA, USA) for data normalization, visualization and graph generation.

### Bio-Layer Interferometry

BLI was performed using single-use Anti-Penta-HIS (HIS1K) sensors on an eight-channel Octet RED 96e instrument according to manufacturer’s protocols (Fortebio). Assays were performed in 50 mM Hepes pH 8.0, 150 mM NaCl, supplemented with 0.05% Tween-20 and 1% BSA to reduce non-specific binding.

For kinetics experiments, the eight sensors were divided into two sets, samples and reference. The sample sensors were loaded with Nb (immobilization level of 0.2-0.25 nm) while reference sensors were not. Titrations were measured as sample/reference pairs at identical H3 concentrations. H3 was used at concentrations ranging from 10 nM to 1000 nM. For comparative binding test, seven sample sensors were loaded with different Nbs, and the remaining sensor was used as reference. Sensors were all dipped into H3 at a concentration of 1µM. Final presented curves were reference-subtracted Savitzky-Golay filtered, as implemented in the Octet® Analysis Studio Software (Sartorius). Global Bayesan 1:1 model fits were obtained using Evilfit (65). GraphPad Prism 9.0 software (CA, USA) was used for visualization and graph generation.

### Isothermal Titration Calorimetry

For ITC, both H3 and Nbs were buffer exchanged to 50 mM Hepes, 150 mM NaCl, pH 8.0 and isolated as single peak population by Superdex 200 increase 10/600 column. ITC measurements were performed using an microcal PEAQ ITC calorimeter (Malvern). The cell temperature was set to 37 °C and the syringe stirring speed to 750 rpm. H3 was loaded into the cell at a concentration of 10 µM, whereas the Nbs were loaded into the syringe at concentrations ranging from 40 to 100 μM. Nb to buffer titrations were performed as control. Shown data were only baseline-corrected, since dilution effects upon titration were not evident. Data and binding parameters were analyzed using the MicroCal Peak ITC software (Malvern). For each H3-Nb, integration and global analysis comprising at least two experiments were performed using software packages NITPIC and SEDPHAT as described in (66). Raw and processed ITC data were imported into GraphPad Prism 9.0 software (CA, USA) for visualization and graphs generation.

### Crystallization and structures determination

Crystallization experiments were performed at 293 K using the sitting drop vapor diffusion method by mixing an equal amount H3-Nb complex and reservoir solution. The initial concentration of H3-Nb complexes were 11 mg/mL and 15 mg/mL for H3-C4 and H3-B5, respectively. Best diffracting crystals were obtained in 0.1 M Sodium citrate, 20% w/v PEG 4000, pH 4.5 for H3-C4; 1.5 M Ammonium sulfate, 0.1M Tris-HCl pH 8.0 for H3-B5. Crystals were cryoprotected with 25% glycerol and flash-frozen in liquid nitrogen. X-ray diffraction data were collected at the beam line ID30B at ESRF (Grenoble, France). Diffraction data of H3-C4 were processed using Staraniso (67), and intensities were merged with AIMLESS (68); for H3-B5 AutoPROC/staraniso autoprocessed data (67) were used and intensities merged with AIMLESS (68). The crystal structures of H3-C4 was determined by molecular replacement using several consecutive runs of PhaserMR (69). Firstly, the H3 constant domain from (27) was used as searching model, followed by a second run searching for the variable and, eventually, a third run searching for the homology model of C4 obtained by Swiss model (70). The crystal structure of H3-B5 were determined by molecular replacement using PhaserMR (69) and the previously determined structure of H3-C4 as searching model. The asymmetric unit was differently organized in the two complexes. In H3-C4 a monomeric H3-C4 complex molecule was present, while in H3-B5 a dimeric H3 was decorated by two B5 molecules. The molecular models were preliminary subjected to rigid-body refinement, followed by restrained refinement using phenix.phaser (71). Manual model building was thereafter carried out using COOT (72).

The energetics involved in complex formation were computed by PDBePISA software (73). Models for H3-H19, H3-C10, and H3-C2 were obtained from Alphafold2 (74). Interactions within H3 and H3-Nb complexes were computed using the PLIP software (75).

### Statistical analysis

*C. elegans* experiments were performed using 100 worms per group and were repeated at least three times based on methods described on http://www.wormbook.org. No randomization was required for *C. elegans* experiments. All evaluations were done blind to the treatment group and sample identity. The data were analyzed using GraphPad Prism 9.0 software (CA, USA) 9.0 software (CA, USA) by one-way or two-way ANOVA, and Bonferroni’s *post hoc* test. A p-value <0.05 was considered significant.

## ACCESSION NUMBER

H3-C4 and H3-B5 atomic coordinates and the structure factors have been deposited in the Protein Data Bank with the following accession numbers 8P88 and 8P89, respectively.

## ACKNOWLEDGEMENTS

We would like to thank Dr. Alain Roussel from NabGen Technology for his kind help and support during the generation and identification of Nbs. This research was funded by the Italian Ministry of Research PRIN 2020 (20207XLJB2) and by Ricerca Corrente from Italian Ministry of Health to IRCCS Policlinico San Donato.

## REFERENCES

1. Merlini G, Dispenzieri A, Sanchorawala V, Schönland SO, Palladini G, Hawkins PN, et al. (2018). Systemic immunoglobulin light chain amyloidosis. Nat Rev Dis Prim. 4(1):38.

2. Merlini G. (2017) AL amyloidosis: from molecular mechanisms to targeted therapies. Hematol Am Soc Hematol Educ Progr. 2017(1):1–12.

3. Merlini G, Palladini G. (2013) Light chain amyloidosis: the heart of the problem. Vol. 98, Haematologica. Italy; p. 1492–5.

4. Grogan M, Dispenzieri A, Gertz MA. (2017) Light-chain cardiac amyloidosis: strategies to promote early diagnosis and cardiac response. Heart. 103(14):1065–72.

5. Palladini G, Milani P, Merlini G. (2020). Management of AL amyloidosis in 2020. Blood. 136(23):2620–7.

6. Palladini G, Dispenzieri A, Gertz MA, Kumar S, Wechalekar A, Hawkins PN, et al. (2012). New criteria for response to treatment in immunoglobulin light chain amyloidosis based on free light chain measurement and cardiac biomarkers: impact on survival outcomes. J. Am. Soc. Clin. Oncol. 30(36):4541–9.

7. Lavatelli F. (2022). Mechanisms of Organ Damage and Novel Treatment Targets in AL Amyloidosis. Vol. 3, Hemato. p. 47–62.

8. Maritan M, Romeo M, Oberti L, Sormanni P, Tasaki M, Russo R, et al. (2020). Inherent Biophysical Properties Modulate the Toxicity of Soluble Amyloidogenic Light Chains. J Mol Biol. 432(4):845–60.

9. Shi J, Guan J, Jiang B, Brenner DA, del Monte F, Ward JE, et al. (2010). Amyloidogenic light chains induce cardiomyocyte contractile dysfunction and apoptosis via a non-canonical p38α MAPK pathway. Proc Natl Acad Sci. 107(9):4188–93.

10. Marin-Argany M, Lin Y, Misra P, Williams A, Wall JS, Howell KG, et al. (2016). Cell Damage in Light Chain Amyloidosis: fibril internalization, toxicity, and cell-mediated seeding. J Biol Chem. 291(38):19813–25.

11. Brenner DA, Jain M, Pimentel DR, Wang B, Connors LH, Skinner M, et al. (2004) Human Amyloidogenic Light Chains Directly Impair Cardiomyocyte Function Through an Increase in Cellular Oxidant Stress. Circ Res. 94(8):1008–10.

12. Marin-Argany M, Güell-Bosch J, Blancas-Mejía LM, Villegas S, Ramirez-Alvarado M. (2015). Mutations can cause light chains to be too stable or too unstable to form amyloid fibrils. Protein Sci. 24(11):1829–40.

13. Imperlini E, Gnecchi M, Rognoni P, Sabidò E, Ciuffreda MC, Palladini G, et al. (2017). Proteotoxicity in cardiac amyloidosis: amyloidogenic light chains affect the levels of intracellular proteins in human heart cells. Sci Rep. 7(1):15661.

14. Lavatelli F, Imperiini E, Orrù S, Rognoni P, Sarnataro D, Palladini G, et al. (2015). Novel mitochondrial protein interactors of immunoglobulin light chains causing heart amyloidosis. FASEB J. 29(11):4614–28.

15. Palladini G, Lavatelli F, Russo P, Perlini S, Perfetti V, Bosoni T, et al. (2006). Circulating amyloidogenic free light chains and serum N-terminal natriuretic peptide type B decrease simultaneously in association with improvement of survival in AL. Blood. 107(10):3854–8.

16. Palladini G, Milani P, Merlini G. (2022). In search of the most effective therapy for light chain amyloidosis. Amyloid. 29(1):67–8.

17. McWilliams-Koeppen HP, Foster JS, Hackenbrack N, Ramirez-Alvarado M, Donohoe D, Williams A, et al. (2015) Light Chain Amyloid Fibrils Cause Metabolic Dysfunction in Human Cardiomyocytes. PLoS One. 10(9):e0137716.

18. Levinson RT, Olatoye OO, Randles EG, Howell KG, DiCostanzo AC, Ramirez-Alvarado M. (2013). Role of mutations in the cellular internalization of amyloidogenic light chains into cardiomyocytes. Sci Rep. 3:1278.

19. Sikkink LA, Ramirez-Alvarado M. (2010) Cytotoxicity of amyloidogenic immunoglobulin light chains in cell culture. Cell Death Dis. 1(11):98.

20. Shi J, Guan J, Jiang B, Brenner DA, Del Monte F, Ward JE, et al. (2010). Amyloidogenic light chains induce cardiomyocyte contractile dysfunction and apoptosis via a non-canonical p38alpha MAPK pathway. Proc Natl Acad Sci. 107(9):4188–93.

21. Liao R, Jain M, Teller P, Connors LH, Ngoy S, Skinner M, et al. (2001). Infusion of light chains from patients with cardiac amyloidosis causes diastolic dysfunction in isolated mouse hearts. Circulation. 104(14):1594–7.

22. Shin JT, Ward JE, Collins PA, Dai M, Semigran HL, Semigran MJ, et al. (2012) Overexpression of human amyloidogenic light chains causes heart failure in embryonic zebrafish: a preliminary report. Amyloid. 19(4):191–6.

23. Mishra S, Guan J, Plovie E, Seldin DC, Connors LH, Merlini G, et al. (2013). Human amyloidogenic light chain proteins result in cardiac dysfunction, cell death, and early mortality in zebrafish. Am J Physiol Heart Circ Physiol. 305(1):H95–103.

24. Diomede L, Rognoni P, Lavatelli F, Romeo M, del Favero E, Cantù L, et al. (2014). A Caenorhabditis elegans-based assay recognizes immunoglobulin light chains causing heart amyloidosis. Blood. 123(23):3543–52.

25. Diomede L, Romeo M, Rognoni P, Beeg M, Foray C, Ghibaudi E, et al. (2017). Cardiac Light Chain Amyloidosis: The Role of Metal Ions in Oxidative Stress and Mitochondrial Damage. Antioxid Redox Signal. 27(9):567–82.

26. Russo R, Romeo M, Schulte T, Maritan M, Oberti L, Barzago MM, et al. (2022). Cu(II) Binding Increases the Soluble Toxicity of Amyloidogenic Light Chains. Int J Mol Sci. 23(2).

27. Oberti L, Rognoni P, Barbiroli A, Lavatelli F, Russo R, Maritan M, et al. (2017). Concurrent structural and biophysical traits link with immunoglobulin light chains amyloid propensity. Sci Rep.7(1):16809.

28. Absmeier RM, Rottenaicher GJ, Svilenov HL, Kazman P, Buchner J. (2022). Antibodies gone bad – the molecular mechanism of light chain amyloidosis. FEBS J.

29. Weber B, Hora M, Kazman P, Pradhan T, Rührnößl F, Reif B, et al. (2020). Domain Interactions Determine the Amyloidogenicity of Antibody Light Chain Mutants. J Mol Biol. 432(23):6187–99.

30. Blancas-Mejía LM, Horn TJ, Marin-Argany M, Auton M, Tischer A, Ramirez-Alvarado M. (2015). Thermodynamic and fibril formation studies of full length immunoglobulin light chain AL-09 and its germline protein using scan rate dependent thermal unfolding. Biophys Chem. 207:13–20.

31. del Pozo-Yauner L, Wall JS, González Andrade M, Sánchez-López R, Rodríguez-Ambriz SL, Pérez Carreón JI, et al. (2014). The N-terminal strand modulates immunoglobulin light chain fibrillogenesis. Biochem Biophys Res Commun. 443(2):495–9.

32. Rennella E, Morgan GJ, Yan N, Kelly JW, Kay LE. (2019). The Role of Protein Thermodynamics and Primary Structure in Fibrillogenesis of Variable Domains from Immunoglobulin Light Chains. J Am Chem Soc. 141(34):13562–71.

33. Morgan GJ, Kelly JW. (2016). The Kinetic Stability of a Full-Length Antibody Light Chain Dimer Determines whether Endoproteolysis Can Release Amyloidogenic Variable Domains. J Mol Biol. 428(21):4280–97.

34. Klimtchuk ES, Gursky O, Patel RS, Laporte KL, Connors LH, Skinner M, et al. (2010). The Critical Role of the Constant Region in Thermal Stability and Aggregation of Amyloidogenic Immunoglobulin Light Chain. Biochemistry. 49(45):9848–57.

35. Rennella E, Morgan GJ, Kelly JW, Kay LE. (2019). Role of domain interactions in the aggregation of full-length immunoglobulin light chains. Proc Natl Acad Sci. 116(3):854– 63.

36. Morgan GJ, Yan NL, Mortenson DE, Rennella E, Blundon JM, Gwin RM, et al. (2019). Stabilization of amyloidogenic immunoglobulin light chains by small molecules. Proc Natl Acad Sci. 116(17):8360–9.

37. Yan NL, Morgan GJ, Petrassi HM, Wilson IA, Kelly JW. (2023). Pharmacological stabilization of the native state of full-length immunoglobulin light chains to treat light chain amyloidosis. Curr Opin Chem Biol. 75:102319.

38. Morgan GJ, Buxbaum JN, Kelly JW. (2021). Light Chain Stabilization: A Therapeutic Approach to Ameliorate AL Amyloidosis. Hemato. 2(4):645–59.

39. Muyldermans S. (2013). Nanobodies: natural single-domain antibodies. Annu Rev Biochem. 82:775–97.

40. Bao G, Tang M, Zhao J, Zhu X. (2021). Nanobody: a promising toolkit for molecular imaging and disease therapy. EJNMMI Res. 11(1):6.

41. Kolkman JA, Law DA. (2010). Nanobodies – from llamas to therapeutic proteins. Drug Discov Today Technol. 7(2):e139–46.

42. Bourne PC, Ramsland PA, Shan L, Fan Z-C, DeWitt CR, Shultz BB, et al. (2002). Three-dimensional structure of an immunoglobulin light-chain dimer with amyloidogenic properties. Acta Crystallogr Sect D. 58(5):815–23.

43. Yan NL, Santos-Martins D, Rennella E, Sanchez BB, Chen JS, Kay LE, et al. (2020). Structural basis for the stabilization of amyloidogenic immunoglobulin light chains by hydantoins. Bioorg Med Chem Lett. 30(16):127356.

44. Brumshtein B, Esswein SR, Salwinski L, Phillips ML, Ly AT, Cascio D, et al. (2015). Inhibition by small-molecule ligands of formation of amyloid fibrils of an immunoglobulin light chain variable domain. Elife. 4:e10935.

45. Brumshtein B, Esswein SR, Landau M, Ryan CM, Whitelegge JP, Phillips ML, et al. (2014). Formation of amyloid fibers by monomeric light chain variable domains. J Biol Chem. 289(40):27513–25.

46. Baden EM, Owen BAL, Peterson FC, Volkman BF, Ramirez-Alvarado M, Thompson JR. (2008). Altered dimer interface decreases stability in an amyloidogenic protein. J Biol Chem. 283(23):15853–60.

47. Radamaker L, Lin Y-H, Annamalai K, Huhn S, Hegenbart U, Schönland SO, et al. (2019). Cryo-EM structure of a light chain-derived amyloid fibril from a patient with systemic AL amyloidosis. Nat Commun. 10(1):1103.

48. Radamaker L, Karimi-Farsijani S, Andreotti G, Baur J, Neumann M, Schreiner S, et al. (2021). Role of mutations and post-translational modifications in systemic AL amyloidosis studied by cryo-EM. Nat Commun. 12(1):6434.

49. Radamaker L, Baur J, Huhn S, Haupt C, Hegenbart U, Schönland S, et al. (2021). Cryo-EM reveals structural breaks in a patient-derived amyloid fibril from systemic AL amyloidosis. Nat Commun. 12(1):875.

50. Swuec P, Lavatelli F, Tasaki M, Paissoni C, Rognoni P, Maritan M, et al. (2019). Cryo-EM structure of cardiac amyloid fibrils from an immunoglobulin light chain AL amyloidosis patient. Nat Commun. 10(1):1269.

51. Fusco G, Chen SW, Williamson PTF, Cascella R, Perni M, Jarvis JA, et al. (2017). Structural basis of membrane disruption and cellular toxicity by α-synuclein oligomers. Science. 358(6369):1440–3.

52. Iyer A, Claessens MMAE. (2019). Disruptive membrane interactions of alpha-synuclein aggregates. Biochim Biophys Acta - Proteins Proteomics. 1867(5):468–82.

53. Eisenberg DS, Sawaya MR. (2017). Structural Studies of Amyloid Proteins at the Molecular Level. Annu Rev Biochem. 86:69–95.

54. Lewkowicz E, Gursky O. (2022). Dynamic protein structures in normal function and pathologic misfolding in systemic amyloidosis. Biophys Chem. 280:106699.

55. Kazman P, Vielberg M-T, Pulido Cendales MD, Hunziger L, Weber B, Hegenbart U, et al. (2020). Fatal amyloid formation in a patient’s antibody light chain is caused by a single point mutation. Elife. 9:e52300.

56. Otzen DE. (2021). Driving forces in amyloidosis: How does a light chain make a heavy heart? J Biol Chem. 296:100785.

57. Rottenaicher GJ, Weber B, Rührnößl F, Kazman P, Absmeier RM, Hitzenberger M, et al. (2021). Molecular mechanism of amyloidogenic mutations in hypervariable regions of antibody light chains. J Biol Chem. 296:100334.

58. Rottenaicher GJ, Absmeier RM, Meier L, Zacharias M, Buchner J. (2023). A constant domain mutation in a patient-derived antibody light chain reveals principles of AL amyloidosis. Commun Biol. 6(1):209.

59. Klimtchuk ES, Peterle D, Bullitt EA, Connors LH, Engen JR, Gursky O. (2023). Role of Complementarity-Determining Regions 1 and 3 in Pathologic Amyloid Formation by Human Immunoglobulin κ1 Light Chains. bioRxiv. 2023.02.01.526662.

60. Lavatelli F, Mazzini G, Ricagno S, Iavarone F, Rognoni P, Milani P, et al. (2020). Mass spectrometry characterization of light chain fragmentation sites in cardiac AL amyloidosis: insights into the timing of proteolysis. J Biol Chem. 295(49):16572–84.

61. Lavatelli F, Perlman DH, Spencer B, Prokaeva T, McComb ME, Théberge R, et al. (2008). Amyloidogenic and associated proteins in systemic amyloidosis proteome of adipose tissue. Mol Cell Proteomics. 7(8):1570–83.

62. Mazzini G, Ricagno S, Caminito S, Rognoni P, Milani P, Nuvolone M, et al. (2022). Protease-sensitive regions in amyloid light chains: what a common pattern of fragmentation across organs suggests about aggregation. FEBS J. 289(2):494–506.

63. Arbabi Ghahroudi M, Desmyter A, Wyns L, Hamers R, Muyldermans S. (1997). Selection and identification of single domain antibody fragments from camel heavy-chain antibodies. FEBS Lett. 414(3):521–6.

64. Desmyter A, Spinelli S, Roussel A, Cambillau C. (2015). Camelid nanobodies: killing two birds with one stone. Curr Opin Struct Biol. 32:1–8.

65. Svitel J, Balbo A, Mariuzza RA, Gonzales NR, Schuck P. (2003). Combined affinity and rate constant distributions of ligand populations from experimental surface binding kinetics and equilibria. Biophys J. 84(6):4062–77.

66. Brautigam CA, Zhao H, Vargas C, Keller S, Schuck P. (2016). Integration and global analysis of isothermal titration calorimetry data for studying macromolecular interactions. Nat Protoc. 11(5):882–94.

67. Vonrhein C, Tickle IJ, Flensburg C, Keller P, Paciorek W, Sharff A, et al. (2018). Advances in automated data analysis and processing within autoPROC, combined with improved characterisation, mitigation and visualisation of the anisotropy of diffraction limits using STARANISO. Acta Crystallogr Sect A. 74, 360.

68. Evans PR, Murshudov GN. (2013). How good are my data and what is the resolution? Acta Crystallogr D Biol Crystallogr. 69:1204–14.

69. McCoy AJ, Grosse-Kunstleve RW, Adams PD, Winn MD, Storoni LC, Read RJ. (2007). Phaser crystallographic software. J Appl Crystallogr. 40:658–74.

70. Waterhouse A, Bertoni M, Bienert S, Studer G, Tauriello G, Gumienny R, et al. (2018). SWISS-MODEL: homology modelling of protein structures and complexes. Nucleic Acids Res. 46:W296–303.

71. Liebschner D, Afonine P V, Baker ML, Bunkóczi G, Chen VB, Croll TI, et al. (2019). Macromolecular structure determination using X-rays, neutrons and electrons: recent developments in Phenix. Acta Crystallogr Sect D, Struct Biol. 75:861–77.

72. Emsley P, Cowtan K. (2004). Coot: model-building tools for molecular graphics. Acta Crystallogr D Biol Crystallogr. 60:2126–32.

73. Krissinel E, Henrick K. (2007). Inference of macromolecular assemblies from crystalline state. J Mol Biol. 372(3):774–97.

74. Mirdita M, Schütze K, Moriwaki Y, Heo L, Ovchinnikov S, Steinegger M. (2022). ColabFold: making protein folding accessible to all. Nat Methods. 19(6):679–82.

75. Salentin S, Schreiber S, Haupt VJ, Adasme MF, Schroeder M. (2015). PLIP: fully automated protein–ligand interaction profiler. Nucleic Acids Res. 43(W1):W443–7.

