## Supplemental data for "Nanobodies counteract the toxicity of an amyloidogenic light chain by stabilizing a partially open dimeric conformation"

### SUPPLEMENTARY TABLES

**Table S1.** Primary sequence and molecular weight of the five llama-derived Nbs targeting H3.

| **Nb** | **Primary sequence** | **MW kDa** |
|---|---|---|
| C2 | QVQLVESGGGSVQPGGSLRLSCAASGFTFSNDDMSWVRQGPGKGLEWVSSIRSGGGATSYSDSVKGRFTISRDDHKNMLYLEMNSL KPDDTAVYYCVKLGVTGWGPPQDDYWGQGTQVTVSS | 13.1 |
| C4 | QVQLVESGGAAVQTGGSLRLSCVASGFDLSNHAMAWVRQSPGKGLEYISGINNGGTTTTYGDSVKDRFTISRDNAKSTVYLQMNRLE ADDTAVYYCVKSSYSDIVSARFDSWGKGTQVTVSS | 13.1 |
| B5 | QVQLVESGGALVEPGGSLRLSCVASGFPFNEYDMSWVRQAPGKGLEWVSTIKSDGAYTYYGDNVKGRFTISRDNAKNTLYLQMNSL KSEDTGVYYCAKEARIGVSGTYDYWGQGTQVTVSS | 13.2 |
| C10 | QVQLVESGGGLVQPGGSLRLSCAASGFTFDDYAMSWVRQAPGKGLEWVSAISWNGGSTFYAESMKGRFTISRDNAKNTLYLQMNG LKSEDTAVYYCAKTNWGHLGVMDSWGKGTQVTVSS | 13.0 |
| H19 | QVQLVESGGGLVQPGGSLRLSCAASGFTFGIYDMSWVRQAPGKGPEWVSTINSGGGSTYYADSVKGRFTISRDNTKNTVYLQMNSL QSEDTALYYCAKEMNGPNRAYDYWGQGTQVTVSS | 13.0 |

**Table S2.** Hydrophobic Interactions stabilizing H3 $V_L$ - B5, H3 $V_L$ - H3 $V_L$, and H3 $V_L$ - C4 interfaces.

| H3-B5 complex | | | H3 homodimer (from PDB 5MTL) | | | H3-C4 complex | | |
|---|---|---|---|---|---|---|---|---|
| Residue H3 | Residue Nb | Distance / Å | Residue H3 | Residue H3 | Distance / Å | Residue H3 | Residue Nb | Distance / Å |
| Phe101 | Val37 | 3.85 | Phe101 | Tyr37 | 3.99 | Phe101 | Leu45 | 4.00 |
| Phe88 | Leu45 | 3.54 | Phe88 | Ala44 | 3.53 | Phe88 | Leu45 | 3.97 |
| Pro45 | Leu45 | 3.85 | Ala44 | Phe88 | 3.78 | Val99 | Tyr47 | 3.71 |
| Val99 | Trp47 | 3.72 | Pro45 | Phe101 | 3.40 | Ala44 | Tyr95 | 3.95 |
| Leu96 | Asp62 | 3.98 | | | | Pro45 | Trp112 | 3.90 |
| Ala44 | Tyr95 | 3.94 | | | | Tyr37 | Trp112 | 3.80 |
| Tyr37 | Trp111 | 3.46 | | | | | | |
| Ala44 | Trp111 | 3.88 | | | | | | |

**Table S3.** H-bond and π stacking Interactions stabilizing H3 $V_L$ - B5, H3 $V_L$ - H3 $V_L$, and H3 $V_L$ - C4 interfaces.

| | H3-B5 complex | | | | H3 homodimer (from PDB 5MTL) | | | | H3-C4 complex | | | |
|---|---|---|---|---|---|---|---|---|---|---|---|---|
| | H3 | B5 | H-A / Å | D-A / Å | H3 | H3 | H-A / Å | D-A / Å | H3 | C4 | H-A / Å | D-A / Å |
| H bonds | Gln39 | Gln39 | 2.28 | 3.09 | Gln39 | Gln39 | 1.95 | 2.91 | Gln39 | Gln39 | 2.00 | 2.98 |
| | Gln39 | Gln39 | 1.66 | 2.64 | Gln39 | Gln39 | 2.78 | 3.56 | Gln39 | Gln39 | 2.33 | 3.31 |
| | Val99 | Trp47 | 2.39 | 3.30 | | | | | Asn97 | Thr59 | 3.35 | 3.76 |
| | Gln39 | Tyr95 | 3.29 | 3.73 | | | | | Tyr37 | Ala107 | 1.56 | 2.47 |
| | Asn35 | Gly106 | 3.48 | 3.90 | | | | | Tyr37 | Phe109 | 1..92 | 2.74 |
| | Asn35 | Gly106 | 2.30 | 2.78 | | | | | Lys46 | Asp110 | 3.02 | 4.03 |
| | Tyr37 | Gly106 | 1.66 | 2.55 | | | | | Lys46 | Trp112 | 3.00 | 3.75 |
| | Tyr37 | Thr107 | 2.09 | 2.85 | | | | | | | | |
| | H3 | B5 | Distance / Å | | | | | | H3 | C4 | Distance / Å | |
| π stacking | Trp92 | Tyr59 | 5.48 | | | | | | His50 | Arg108 | 3.68 | |

H-A: H-bond distance between hydrogen (H) and acceptor (A) atom;
D-A: H-bond distance between donor (D) and acceptor (A) atoms.

#### SUPPLEMENTARY FIGURES

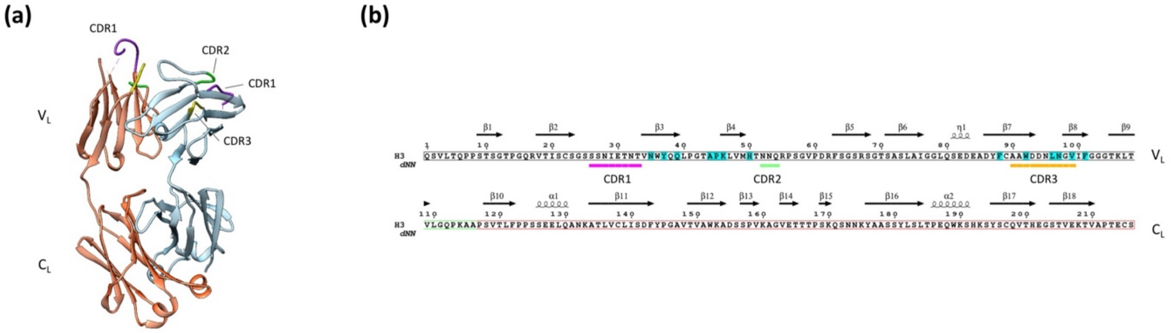


**Figure S1. H3 native structure and sequence.** (a) Homo-dimeric three-dimensional fold of H3 (PDB 5MTL). H3 presents the typical immunoglobulin-like domain of LC, where each LC monomer comprises a N-terminal variable domain ($V_L$) and a C-terminal constant domain ($C_L$). Each $V_L$ domain contains three complementary determining regions (CDR1, purple; CDR2, green; CDR3, yellow). (b) Primary sequence of H3. Residues belonging to the $V_L$ domain are indicated by a black box, while residues of the $C_L$ domain are indicated by a red box. The linker connecting $V_L$ and $C_L$ is indicated by a green box.. Secondary structure elements of H3 are shown above the sequence alignment and were extracted from the H3 structure (PDB 5MTL). CDR1, CDR2, and CDR3 are indicated by pink, green and yellow bars, respectively, positioned below the sequence alignment. Residues involved in H3-Nb interaction are highlighted in cyan. The alignment was visualized using ESPript (41).

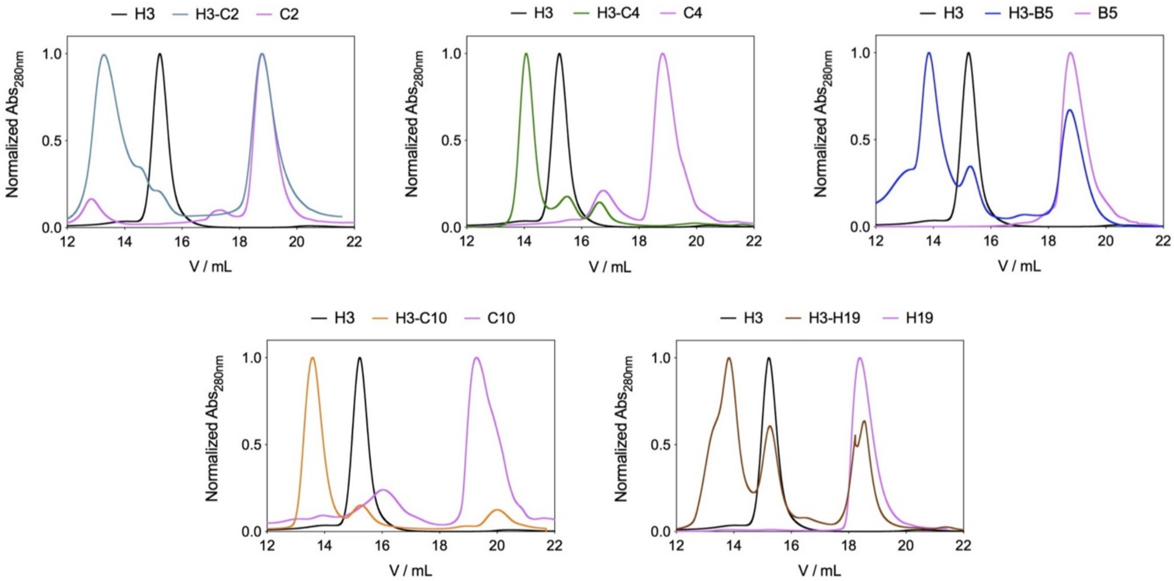


**Figure S2. Analytical size exclusion chromatography analysis.** Size exclusion chromatography analysis of H3, Nb, and H3-Nb complex. H3 is reported as a black line, the Nb as a purple line, and the H3-Nb complex is depicted in a different colour depending on the Nb.

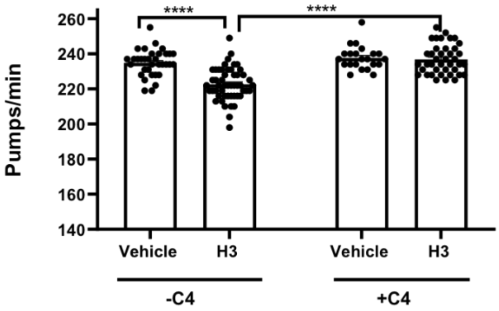


**Figure S3. Nb C4 co-administered with H3, reverted its toxicity in *C. elegans*.** Worms were fed (100 worms/100 µL) with 100 µg/mL H3 (H3 - C4), 100 µg/mL C4 Nb suspended in 10 mM phosphate buffered saline (PBS), pH 7.4 (Vehicle + C4) or 100 µg/mL H3 previously incubated for 30 minutes at room temperature with 100 µg/mL C4 Nb in 10 mM PBS, pH 7.4 (H3 + C4). Control worms were fed 10 mM PBS, pH 7.4 (100 worms/100 µL) (Vehicle – C4). Pharyngeal pumping was determined 24 h after the administration by scoring the number of times the terminal bulb of the pharynx contracted over a 1-min interval (pumps/min). Each value is the mean ± SE (N=50). ****$p<0.0001$, two-way ANOVA and Bonferroni's *post hoc* test. Interaction: C4/H3= ***$p<0.001$, two-way ANOVA and Bonferroni's *post hoc* test.

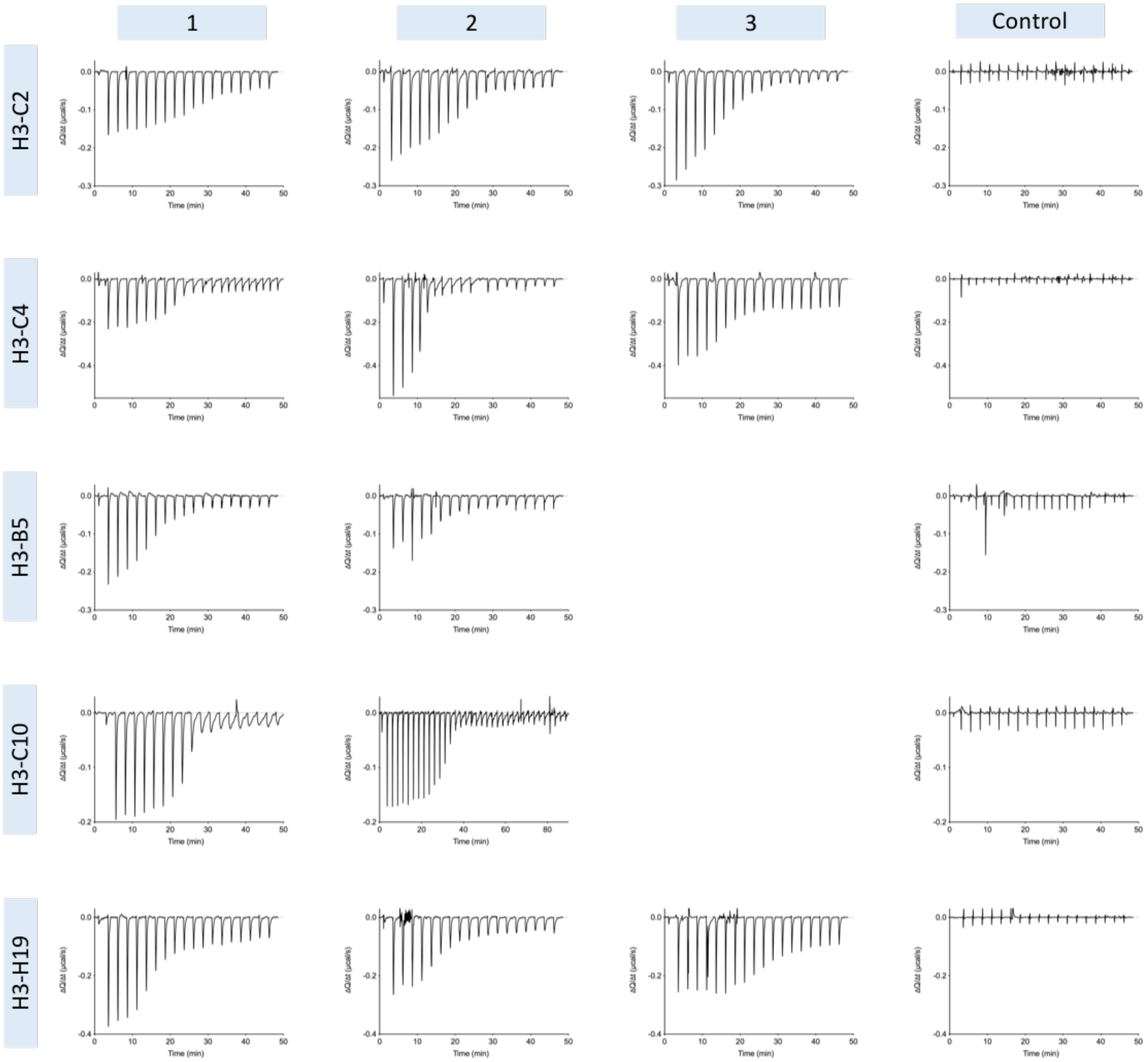


**Figure S4. ITC thermograms.** A complete set of baseline-corrected ITC thermograms used for the ITC binding isotherms is shown in Figure 3A. Plots are organized in rows split by H3/Nb combination and columns split by experiment number. Nb to buffer titrations were performed as control.

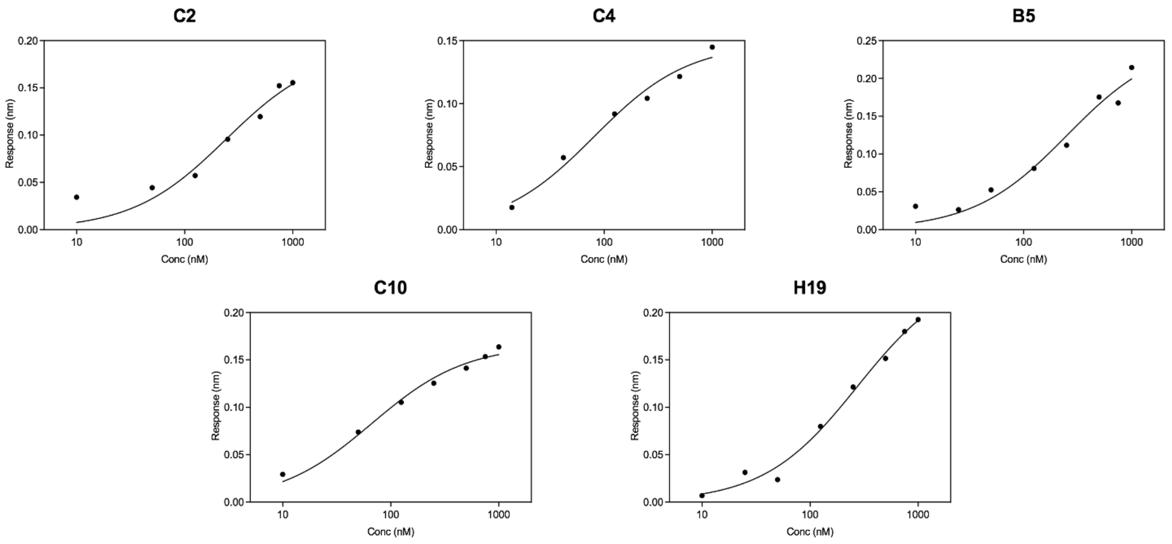


**Figure S5. BLI dose response curves.** Pseudo-equilibrium response values plotted against the logarithmic H3 concentration revealed sigmoidal binding curves that were fit to the single-site interaction model yielding $K_D$-values in the nM range for all Nbs.

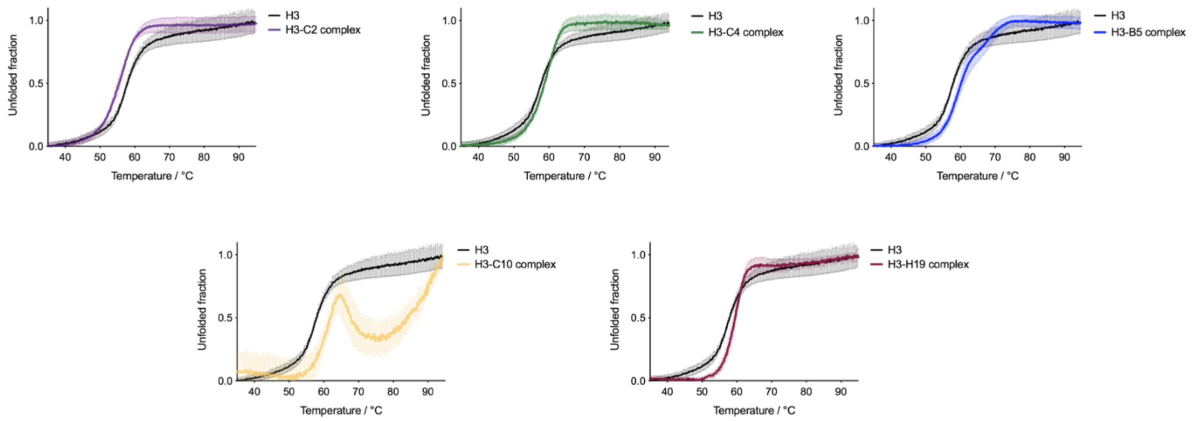


**Figure S6. Thermal unfolding profiles.** Thermal unfolding profiles of H3 (black) and H3-Nb complexes with error bars. Melting temperature is defined as the temperature at which the folding-to-unfolding transition occurs and is the temperature at which the second derivative of the trace is equal to zero.

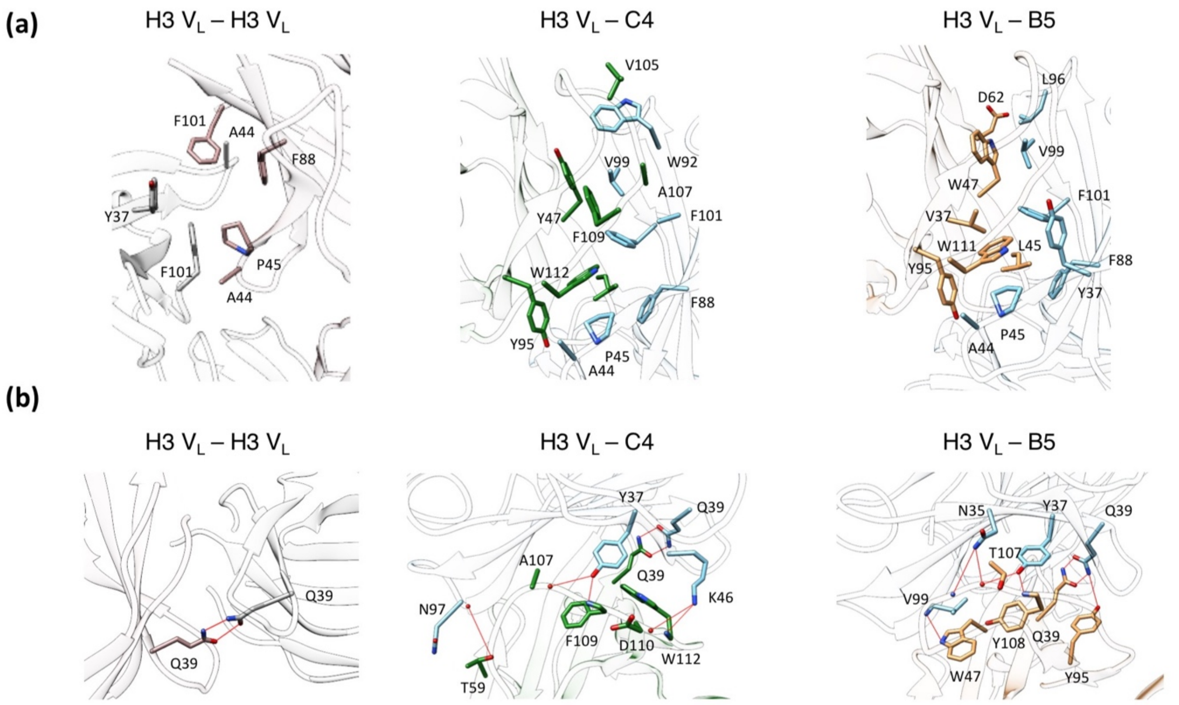


**Figure S7. The H3-Nb interface is tightly stabilized by H-bond and hydrophobic interactions.** (A) Hydrophobic interactions stabilizing the H3 $V_L$ - H3 $V_L$ (left panel), H3 $V_L$ (cyan) – C4 (green) (middle panel), and H3 $V_L$ (cyan) - B5 (orange) interfaces (right panel). Residues involved in hydrophobic interactions are represented as sticks. (b) Ionic and H bond interactions stabilizing the H3 $V_L$ - H3 $V_L$ (left panel), H3 $V_L$ (cyan) – C4 (green) (middle panel), and H3 $V_L$ (cyan) - B5 (orange) interfaces (right panel). Residues involved in H-bond contacts are represented as sticks. H-bond are drawn as red lines. See supplementary table S4 for distances.

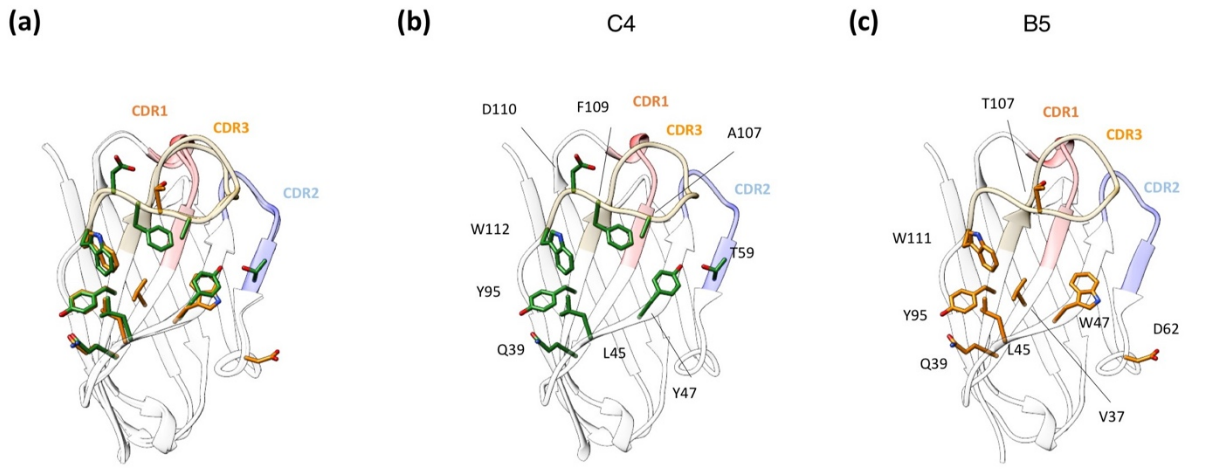


**Figure S8. C4 and B5 share specific positions on their three-dimensional folds to optimize the binding to H3**. (a) Superimposition of Nbs C4 (green) and B5 (orange). Residues involved in Nb-H3 $V_L$ interaction are represented as sticks and color-coded according to Nb. CDR1, CDR2, and CDR3 are depicted in red, blue, and orange, respectively. The positions on Nb fold that are crucial for the interaction with H3 $V_L$ are highly conserved among the two Nbs. In particular, residues Y95, E39, and L45 establish conserved H-bond (Y95 and E39) and hydrophobic interaction (L45) with the target H3 in both complexes. It is worth noticing that the differences in aminoacid composition in crucial positions for the binding are conservative, meaning that the chemistry of the side chains remains mostly unchanged and the interaction network stabilizing H3-Nb interfaces is shared. (b) C4 structure with residues involved in H3 $V_L$ binding represented as sticks. CDRs are reported and highlighted in red (CDR1), blue (CDR2), and orange (CDR3). (c) B5 structure with residues involved in H3 $V_L$ binding represented as sticks. CDRs are reported and highlighted in red (CDR1), blue (CDR2), and orange (CDR3).

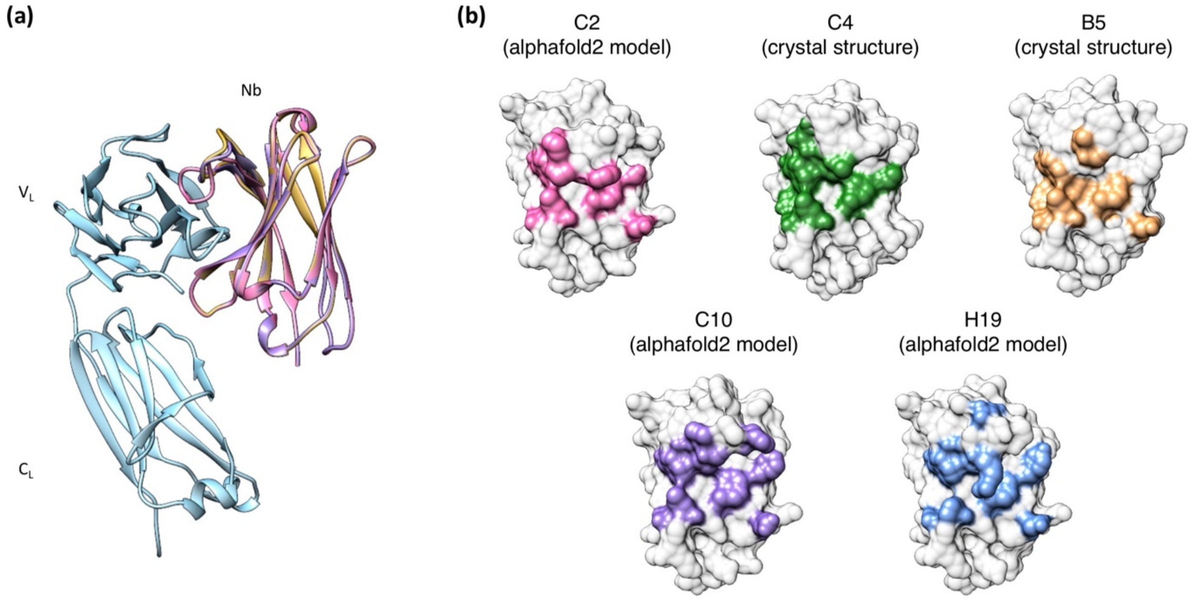


**Figure S9. Nbs C10, C2, and H19 are predicted to engage with H3 in a comparable manner to Nbs C4 and B5.** (a) Superimposition of alphafold2 models of H3-C2 (pink), H3-C10 (purple), and H3-H19 (blue) to crystal structures of H3-C4 (green) and H3-B5 (orange) shows that all Nbs are predicted to bind in an equal manner to H3 (cyan). The alignment was done on the $C_L$ domains, both for crystal structure models and for prediction models. (b) Surface representation of Nbs with residues involved in H3 $V_L$ – Nb interface color-coded according to Nb.

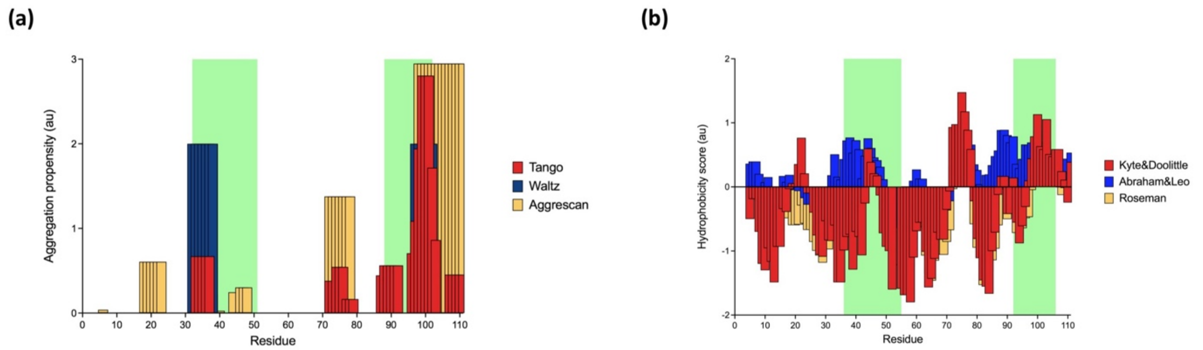


**Figure S10. Nbs interact with aggregation prone and hydrophobic regions of H3 $V_L$** (a) Intrinsic aggregation propensities profiles of H3 $V_L$ computed using three different algorithms, Tango (red) (1), Waltz (Blue) (2), and Aggrescan (Yellow) (3). Each algorithm uses a different scale, the absolute values reported are not intended for comparison among algorithms. Regions of H3 $V_L$ involved in interactions with Nbs are highlighted in green. (b) Hydrophobicity scores of H3 $V_L$ aminoacids were computed using three different algorithms, Kyle&Doolittle (red), Abraham&Leo (Blue), and Rodman (Yellow). Values greater than 0 indicate hydrophobic residues. Regions of H3 $V_L$ involved in interactions with Nbs are highlighted in green.

**REFERENCES**


1. Linding R, Schymkowitz J, Rousseau F, Diella F, Serrano L. (2004). A comparative study of the relationship between protein structure and beta-aggregation in globular and intrinsically disordered proteins. *J Mol Biol*. **342**:345–53.
2. Louros N, Konstantoulea K, De Vleeschouwer M, Ramakers M, Schymkowitz J, Rousseau F. (2020) WALTZ-DB 2.0: an updated database containing structural information of experimentally determined amyloid-forming peptides. *Nucleic Acids Res.* **48**:D389–93.
3. Conchillo-Solé O, de Groot NS, Avilés FX, Vendrell J, Daura X, Ventura S. (2007) AGGRESCAN: a server for the prediction and evaluation of "hot spots" of aggregation in polypeptides. *BMC Bioinformatics*. **8**:65.
